# Whole-organism screening in a zebrafish model of CLN2 disease identifies pregnenolone as a modulator of lysosomal functions with anti-epileptic properties

**DOI:** 10.1101/2025.08.26.670480

**Authors:** Lisa N. Kiani, Gabriele Civiletto, Giulia Lizzo, Anselm Zdebik, Gini Brickell, Fahad Mahmood, Dionysios D. Nalkos, Lucas Michaelides, Philip Eldridge, Emily M. Young, Hazel McPherson, Michelangelo Campanella, Philipp Gut, Claire Russell

## Abstract

Lysosomal storage disorders (LSDs), a group of inherited genetic diseases, are often associated with early-onset neurodegeneration and refractory epileptic seizures. In CLN2 disease, an LSD caused by recessively inherited dysfunction of lysosomal serine protease Tripeptidyl Peptidase 1 (TPP1), lysosomes are functionally impaired through a characteristic accumulation of subcellular materials. Here, we develop and apply a whole-organism screening workflow in *tpp1^-/-^* zebrafish to identify small molecules that suppress epileptic seizures — a hallmark of the human disease — in this model. Among 640 US Food and Drug Administration-approved drugs, pregnenolone, an endogenous precursor for steroid biosynthesis, efficiently suppresses seizures and cell death in *tpp1^-/-^* zebrafish. Using a semi-automated high-content workflow, we further show that pregnenolone normalizes lysosomal architecture in *tpp1^-/-^* zebrafish. Pregnenolone stimulates steroid hormone biosynthesis and related gene expression, which is dysregulated in *tpp1^-/-^* zebrafish. Taken together, *tpp1^-/-^* zebrafish are a suitable model to study CLN2 disease, in which we have identified pregnenolone as a candidate with therapeutic properties.

## Introduction

The Neuronal Ceroid Lipofuscinoses (NCL) are a group of lysosomal storage disorders (LSDs), also known as Batten disease, which collectively constitute the most common cause of inherited paediatric neurodegenerative disorders.^1,2^ There are currently 13 known forms of NCL, each caused by mutation of individual genes. The majority of affected proteins in NCLs localise to the lysosomal lumen or membrane, however others are localised to other cellular compartments, including the endoplasmic reticulum (ER), Golgi, cytosol and plasma membrane.^3^ Despite diversity in the roles of the affected proteins, the 13 currently classified NCLs share a hallmark accumulation of auto-fluorescent lysosomal storage material and common characteristics including progressive neurodegeneration resulting in motor and cognitive decline, retinopathy leading to blindness, myoclonic epilepsy and severely premature death.^2^ The lysosomal storage material in the NCLs predominantly consists of ceroid-lipopigments, as well as subunit c of mitochondrial ATP synthase (SCMAS) or sphingolipid activator proteins A and D.^3,4^

CLN2 disease, otherwise known as the classic Late Infantile form of NCL (cLINCL), is caused by autosomal recessive inherited mutation in the *CLN2* gene, leading to dysfunction of the encoded lysosomal serine protease enzyme Tripeptidyl Peptidase 1 (TPP1).^5,6^ The disease causes cell death in the central nervous system (CNS) and neural retina.^7^ Children with CLN2 disease will typically present at 2–4 years old. Early symptoms can include seizures, language delay, motor dysfunction, behavioural problems and dementia. Progression of the disease leads to rapid motor and cognitive deterioration, progressive visual impairment, the development of refractory seizures and death normally within early adolescence.^8^ Current treatments for NCLs are limited and focus on symptom management.^9,10^ CLN2 disease is currently the only NCL in which there is a clinically approved treatment that has been shown to effectively attenuate disease progression, in the form of enzyme replacement therapy for the brain,^11^ but peripheral disease is likely to emerge in patients receiving enzyme replacement therapy,^12^ which has already been noted in the retina.^13^ Hence, further treatments are still needed. Development of further therapies is reliant on a deeper understanding of the disease pathophysiology, alongside effective animal models to identify and evaluate potential treatment strategies.

In this study we use a zebrafish model of CLN2 disease to perform a drug screen of 640 FDA-approved compounds. The zebrafish model harbours a premature stop codon mutation in exon 3 of the *tpp1* gene, owing to a single T > A point mutation in the *tpp1^sa00^*^11^ allele (*tpp1^sa001^*^1-/-^, herein referred to as *tpp1^-/-^*). The mutation results in deficiency of the Tpp1 enzyme, confirmed by Western blot and an enzyme activity assay.^14^ Previous characterisation of the *tpp1*^-/-^ zebrafish demonstrated their value as a tool for drug discovery; various aspects of the human CLN2 disease are replicated in the *tpp1^-/-^* zebrafish, including storage accumulation and enlarged lysosomes, motor defects and retinal and neuronal degeneration. These disease phenotypes can be observed from 2 days post-fertilisation (dpf) and the zebrafish do not survive past 7 dpf.^14^ Another notable phenotype of *tpp1^-/-^* zebrafish is a period of increased locomotion, indicative of seizures. By using a combination of qualitative and quantitative phenotypes, we performed a relatively high-throughput drug screen. Our drug screen identified the neurosteroid pregnenolone as a potential therapeutic for CLN2 disease.

Here, we show that pregnenolone reduces duration of movement, distance and velocity of seizure-like locomotion activity in *tpp1*^-/-^ zebrafish, demonstrating a rescue of the phenotype. The anti-epileptic effect of pregnenolone was confirmed by electroencephalogram (EEG) measurement of epileptiform activity in the *tpp1*^-/-^ zebrafish. Using live imaging of a novel transgenic zebrafish line expressing a Lamp1 lysosomal reporter, we go on to show that pregnenolone alleviates the lysosomal phenotype in *tpp1*^-/-^ zebrafish. Furthermore, we show that pregnenolone treatment leads to a reduction in cell death in the CNS and enhanced clearance of autophagic material. Finally, given that pregnenolone is a direct metabolite of cholesterol and the precursor of all steroids, we go on to investigate the cholesterol-hormone pathway in *tpp1*^-/-^ zebrafish. Measurement of steroid hormone levels and whole genome RNA sequencing reveal dysregulation of hormone biosynthesis and related genes in *tpp1*^-/-^ zebrafish compared with their healthy (phenotypically wild-type) ‘WT’ siblings. This observation indicates a possible perturbation in the cholesterol pathway in CLN2 disease pathogenesis, highlighting an important area for further research.

## Materials and methods

### Generation and maintenance of zebrafish

This study is reported in line with the <u>ARRIVE guidelines</u>. Adult zebrafish were maintained in a multi-rack aquarium system and kept on a constant 14/10 hr light/dark cycle at 27–29 °C and pH 6.0–7.5.^15^ Adult zebrafish were fed 2-3 times per day a diet of Hikari, krill and live brineshrimp. Experiments conducted at the Royal Veterinary College were done so with approval from the college, the UK Home Office (PPL: PEB686695, PIL: I5DAE98B4) and local Animal Welfare Ethical Review Board (AWERB) under the Animal (Scientific Procedures) Act 1986 Amendment Regulations 2012. Experiments conducted at Nestlé Research (Lausanne, Switzerland) were done so according to Swiss and European Union ethical guidelines, with approval by the animal experimentation ethical committee of Canton of Vaud (permits VD-H13 and VD3177). Where required, embryos were mechanically dechorionated with forceps. For Schedule 1 culling, adult and embryonic/larval zebrafish were exposed to 0.15 % 2-phenoxyethanol for 10 minutes or 24 hr respectively. Adult zebrafish were then decapitated and embryos/larvae were crushed.

### Generation of *tpp1*^sa0011^ homozygotes and normal healthy siblings (WT)

Zebrafish carrying a T/A point mutation in the *tpp1*^sa0011^ allele were generated by N-ethyl-N-nitrosourea mutagenesis in the Tübingen strain and originally obtained from the Sanger Institute (Cambridge, UK: https://www.sanger.ac.uk/Projects/D_rerio/zmp/) as an outcross to WT Tupfel long fin fish^14^ and subsequently maintained for 12 years by backcrossing to WT Tupfel long fin fish. Adult heterozygous carriers of the *tpp1*^sa0011^ mutation (herein referred to as *tpp1^+/-^*) were identified by KASP genotyping. Briefly, this method involves allelic discrimination determined by qPCR reaction ratio of fluorescent cassettes, FAM and HEX, which correspond to T/A allele-specific primers developed by LGC Biosearch Technologies (https://biosearch-cdn.azureedge.net/assetsv6/KASP-genotyping-chemistry-User-guide.pdf). Embryos for experiments were generated by in-cross of *tpp1^+/-^* zebrafish and raised at 28 °C under standard husbandry conditions. Homozygous embryonic genotypes were assigned owing to the presence of a retinal phenotype on or after 2 dpf.^14^ Normal (WT) siblings were used as controls. Prior to any experiment, zebrafish embryos/larvae were viewed under a dissecting microscope to remove unfertilised embryos and to ensure they were healthy, or in the case of *tpp1*^-/-^, that they did not have any abnormal phenotypes beyond the expected ones (i.e. reduced retina size, small head, small and curved body, heart oedema, seizure-like locomotion). If any other phenotypes or injuries were observed in either WT or *tpp1*^-/-^ zebrafish, they were excluded from the study.

### Generation of transgenic zebrafish and confocal imaging

#### Generation of lines

Transgenic zebrafish *Tg(actc1b:lamp1-ZsGreen)^nei08^*, *Tg(actc1b:lc3-ZsGreen)^nei08^*, T*g(actc1b:tfeb-ZsGreen)^nei08^*and *Tg(actc1b:nls-mCherry)^nei09^*^16^ were independently generated using I-SCEI meganuclease-mediated transgenic insertion into 1-cell stage embryos with *tpp1*^+/-^ background as previously described.^17^ Individual founders were selected as ZsGreen-positive or mCherry-positive for propagation of each transgenic line. For ZsGreen-LC3 and ZsGreen-Lamp1 experiments, subsequent generations were crossed with non-transgenic *tpp1*^+/-^ and the progeny of the second generation were selected for fluorescence and used for imaging experiments. For ZsGreen-TFEB experiments, subsequent generations of T*g(actc1b:tfeb-ZsGreen)^nei08^* and *Tg(actc1b:nls-mCherry)^nei09^* were crossed to each other and the progeny were selected for both ZsGreen and mCherry fluorescence and used for imaging experiments.

#### Image acquisition and analysis

Dechorionated embryos, anaesthetised with 0.016 % tricaine (made in aquarium water, buffered with 4 % 1 M Tris pH 9 and adjusted to pH 7 with NaOH), were mounted laterally on 1.5mm glass coverslips with 1 % low-melting point agarose gel in aquarium water. Live imaging was conducted using a 40x water objective on either a Leica^TM^ SP5 or SP8 confocal microscope.

##### ZsGreen-LC3 and ZsGreen-Lamp1

Confocal images were analysed as maximum projections of 8–10 z-stack images using FIJI software. Binary images were generated by applying a threshold to highlight fluorescent signal and eliminate background; the threshold value was the mean fluorescent intensity of all fish in one experiment multiplied by 1.5 for ZsGreen-LC3 and 2 for ZsGreen-Lamp1. Region of interest was defined manually using the freehand drawing tool or by applying a threshold to outline the total area of fish muscle. Measurements of ZsGreen-Lamp1 were derived using the ‘Analyse Particles’ feature on FIJI. ZsGreen-LC3 was measured as a total area of that identified by threshold. Area and puncta count measurements were normalised to the total area of fish muscle per image.

##### ZsGreen-TFEB and mCherry-nls

Confocal images consisting of 6 z-stack images were acquired and analysed individually. Nuclei were identified by a binary image generated by applying a threshold to highlight fluorescent signal of mCherry-nls and eliminate background; the threshold value was kept consistent per experiment. Region of interest was identified by applying a threshold using the ZsGreen-TFEB channel to find the total area of fish muscle. ZsGreen-TFEB was measured as total raw fluorescent units over raw fluorescent units within the identified nuclei. Area and puncta count measurements were normalised to the total area of fish muscle per image.

### Drug screen

At 48 hours post-fertilisation (hpf), dechorionated *tpp1^-/-^*and WT siblings were placed one embryo per well of a flat bottomed 96 round well plate in 49.5 µl aquarium water as shown (Fig 1). Each plate contained one set of WT + DMSO, one set of *tpp1^-/-^* + Dimethyl sulfoxide (DMSO) and the rest of the plates were sets of *tpp1^-/-^*+ each compound. 0.5 µl of DMSO or compound (Enzo Life Sciences FDA-approved library of known bioactives; 400 µg/ml in DMSO) was added to each well to a final compound concentration of 4 µg/ml. In Screen 1, two *tpp1^-/-^* zebrafish were used per compound and plates were incubated until the fish were 120 hpf and then placed in the DanioVision chamber (Noldus) for a 20 minute recording. Outputs from EthoVision tracking software (Noldus) for each fish were distanced moved, mean velocity, max velocity, time spent moving and number of movement bouts (start velocity 0.4 mm/s, stop velocity 0.2 mm/s). Fish were then qualitatively scored for morphology, and quantitatively survival and touch response before culling. In Screen 2, the number of *tpp1^-/-^* zebrafish per compound was increased to three to improve reliability, and the assays were expanded to gain richer data and to include seizure-like locomotion which can only be detected at 72 hpf as loss of locomotion at later stages means that seizures can no longer be detected through assaying movement. We also improved welfare by reducing the age of zebrafish at the last assay time point. Plates were incubated at 28 °C until scored qualitatively for retinal size and quantitatively for survival at 54 hpf, 73 hpf and 79 hpf and then incubated at 22 °C until scoring for retinal size and survival at 97 hpf, and then culled. They were also assessed quantitatively for seizure-like locomotion using DanioVision for 1 hour (with temperature controlled to 25 °C and the light on) at 72 hpf (total distance moved, time spent moving and number of movement bouts; start velocity 4.5 mm/s, stop velocity 2 mm/s) and for loss of locomotion for 20 minutes at 28 °C at 96 hpf (total distance moved, time spent moving and number of movement bouts; start velocity 0.4 mm/s, stop velocity 0.2 mm/s). Compounds that appeared to improve at least one phenotype were re-tested on 10 animals per group (*tpp1^-/-^* + DMSO, *tpp1^-/-^* + compound, WT + DMSO) using the same screen protocol but compound purchased from a new source. The compound that still had beneficial effects after re-screen (pregnenolone) was retested blind in two replicate experiments following the screen protocol except using approx. Ten zebrafish per group (*tpp1^-/-^* + DMSO, *tpp1^-/-^* + pregnenolone, WT + DMSO, WT + pregnenolone).

**Figure 1:**
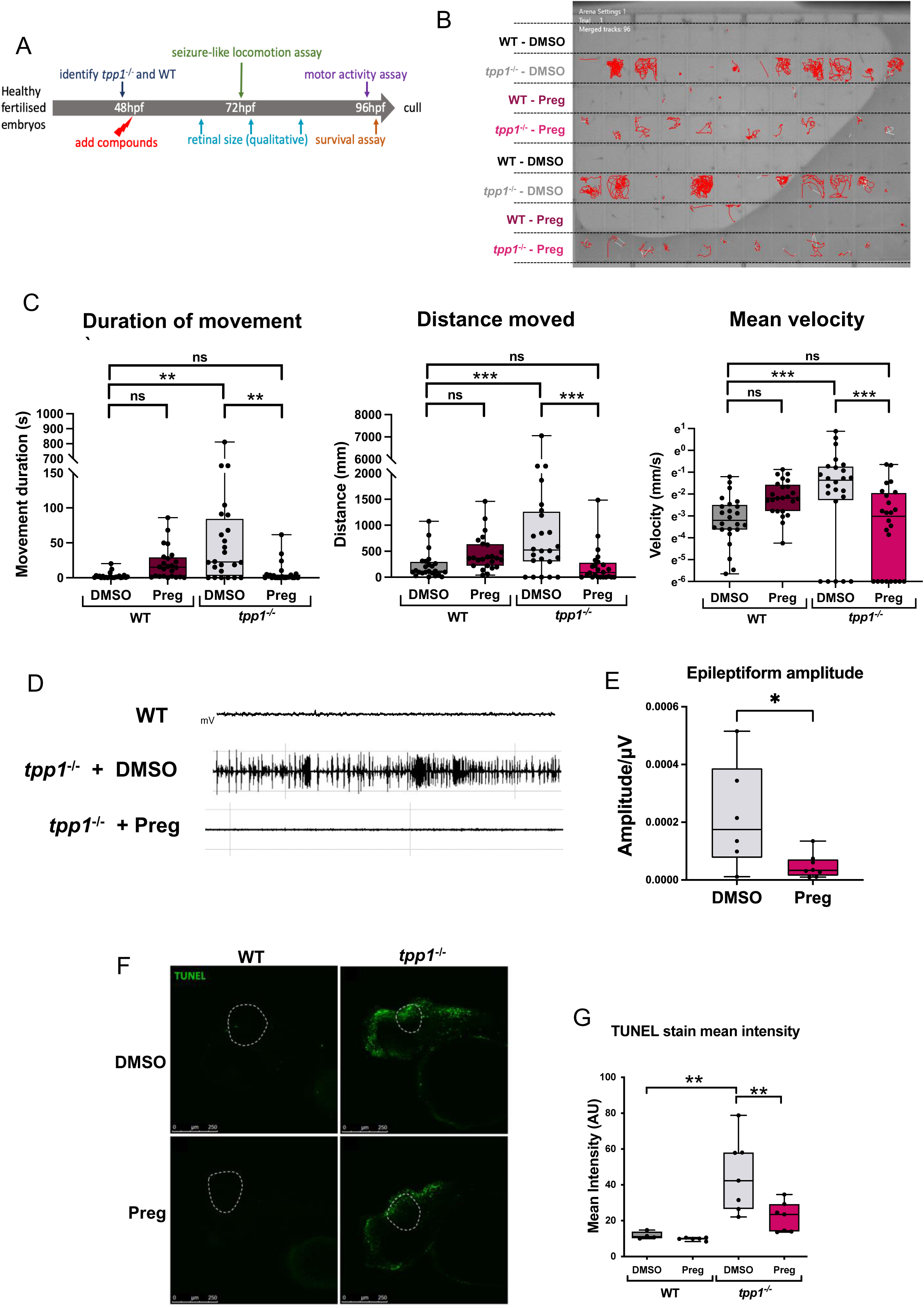
Locomotion assay drug screen in *tpp1^-/-^* zebrafish identifies pregnenolone as hit compound. (A) Drug screen design for FDA approved drug library compounds. (B) Representative locomotion assay plate. 96-well plate containing one fish per well in 200µL aquarium water with treatment. Locomotion tracks are shown in red. White tracks are predicted movements that were not measured. (C) Locomotion assay plots from blind testing of re-sourced pregnenolone in *tpp1^-/-^* zebrafish compared to WT siblings. Pregnenolone (preg) treatment for 24 h from 48 hpf significantly improved seizure-like locomotion phenotype in *tpp1^-/-^* zebrafish, demonstrated by reduced movement duration (s), distance (mm) and mean velocity (mm/s) [Log_Ln_] compared to DMSO treated *tpp1^-/-^*zebrafish. Results from two replicate experiments. (D) Single representative EEG recordings of *tpp1^-/-^* zebrafish at 3 dpf treated with preg or DMSO control for 24 hr. (E) Plot showing significant reduction in mean EEG amplitude in preg treated *tpp1^-/-^* zebrafish compared to DMSO control. Results are combined from three days of experiments. (C and E) Data are shown by box and whisker plot (minimum to maximum) with individual points each representing an individual animal. (C) Linear mixed model (fixed: genotype, treatment / random: date of experiment). (E) Unpaired t-test. *P ≤ 0.033, **P ≤ 0.002, ***P ≤ 0.001. (F) Representative TUNEL stained sagittal sections from WT and *tpp1^-/-^* zebrafish treated with preg or DMSO control from 52 hpf for 3 hours, with the mesencephalon indicated as the region within the dotted white line. (G) Quantification of mean fluorescent intensity of TUNEL staining in the mesencephalon. Results from two replicate experiments. Data are shown by box and whisker plot (minimum to maximum) with individual points each representing an individual fish. Two-way ANOVA. \**p* ≤ 0.033, \*\**p* ≤ 0.002, \*\*\**p* ≤ 0.001.

### Chemical treatments

Embryos were dechorionated mechanically with forceps prior to any treatment.

Pregnenolone (Preg, Sigma, P9129) treatment was performed at 2 dpf for 24–48 hr, as indicated. 0.4 mg/mL pregnenolone dissolved in DMSO was added to aquarium water for a final concentration of 4 μg/mL in 6 well plates with 4.5 mL total volume. 1 % DMSO was used as a control. Dechorionated embryos were incubated in treatment for 24 hr at 28 °C. For 48 hr treatment, treatments were refreshed at 24 hr.

3-Methyladenine (3-MA, Sigma, M9281) treatment^18^ was performed at 2 dpf for 24 hr, alongside or in combination with pregnenolone as above. 3-MA was dissolved in ddH_2_O with gentle heating for a stock concentration of 0.33 M, then added to aquarium water for a final concentration of 10 mM in 6 well plates with 4.5 mL total volume. 1 % DMSO + ddH_2_O was used as a control.

For LC3-ZsGreen flux assay, 500 μL 1 M NH_4_Cl was added directly into 6 well plate containing 4.5 mL of 4 μg/mL pregnenolone (or DMSO control), for a final concentration of 100 mM. The NH_4_Cl was added in the last 4 hr of pregnenolone treatment prior to imaging. 10 % ddH_2_O was used as a control.

### Electroencephalogram (EEG) measurements

The method was adapted from^19^. Following pregnenolone treatment at 2 dpf, 3 dpf larvae were placed in 5mM of D-tubocurarine (Fluka) for 10 minutes, rinsed in aquarium water, and then stabilised in 1.8 % low-melting point agarose gel in aquarium water before placing an electrode onto the optic tectum of the midbrain. Electroencephalogram (EEG) measurements were then recorded for an average of 45 minutes using Axioscope software. Recordings were generated using Clampfit 10 and analysed using Origin 7 software to obtain the mean amplitude.

#### Immunoblotting

Dechorionated 4 dpf larvae maintained on ice, were deyolked by repeat pipetting in Phosphate Buffered Saline (PBS) with 0.1 mM EDTA through a p200 pipette. Samples were then homogenized in RIPA buffer (Sigma) containing 1x protease and phosphatase inhibitors (ThermoScientific), then centrifuged at 5,000 g for 15 minutes at 4 °C to collect the soluble fraction. Protein concentrations were quantified by BCA protein assay (ThermoScientific) and equal concentrations of lysates were mixed with NuPAGE LDS Sample Buffer (4x) (Invitrogen) and NuPAGE Sample Reducing Agent (10x) (Invitrogen), then heated at 70 °C for 10 minutes. Samples were resolved on 4–12 % NuPAGE BiS-Tris gels (Invitrogen) and transferred to nitrocellulose membranes (Invitrogen), which were blocked with 5 % milk powder in TBS-Tween. The membrane was probed with primary antibodies (anti-HSC70 [SantaCruz, sc-7298]; anti-Tom20 [SantaCruz, sc-11415; anti-Subunit C of ATP Synthase (SCMAS) [Abcam, ab110273], anti-P62 [Cell Signalling, 5114S]) overnight at 4 °C and secondary antibodies (anti-mouse/rabbit IgG, HRP-labelled [Perkin Elmer, NEF822001EA/NEF812001EA]) for 1 hr at room temperature, with washing in TBS-Tween in between. Following washing, blots were visualised by ECL kit (ThermoScientific) as directed by manufacturer’s instructions.

#### TUNEL assay

Apoptosis was detected by the DeadEnd^TM^ Fluorometric TUNEL system (Promega), performed based on principles described by Gavrieli *et al.*^20^ using a protocol developed from Promega (2009). Larvae were maintained in 0.003 % 1-phenyl 2-thiourea (PTU) after 24 hpf. Following pregnenolone treatment larvae at 48 hpf, 72 hpf zebrafish were fixed in 4 % paraformaldehyde in PBS (PFA), and stored overnight at 4 °C, before storage in methanol at - 20 °C. Larvae were rehydrated in PBT (50:50 % methanol/PBS solution containing 0.1 % Triton^TM^ X-100 (Sigma-Aldrich)) prior to permeabilization with 10 μg/mL Proteinase K (Promega) in PBT for 60 minutes. Larvae were then washed in PBT, fixed for 20 minutes in 4 % PFA, then chilled for 10 minutes at -20 °C in a 2:1 solution of ethanol and acetone. The larvae were then washed again in PBT before incubation in 100 μl Equilibration Buffer (Promega) on a rocker for 30 minutes, before incubation in TUNEL stain buffer (35 μL Equilibration Buffer, 10 μL Nucleotide Mix, 2 μL Terminal Deoxynucleotidyl Transferase, Recombinant enzyme (Promega) at 37 °C for 60 minutes inside a humified chamber, protected from light. The reaction was terminated by the addition of 500 μl of 2 x SSC (Promega) for 15 minutes, followed by washes in PBT, after which the larvae were cleared in 70 % glycerol/PBS solution. Larvae were mounted laterally on 1.5mm glass coverslips with 2 % low-melting point agarose gel in aquarium water. TUNEL staining was imaged using a Leica SP5 confocal microscope at 10x magnification. Fluorescent intensity was quantified using Volocity software version 6.3 (PerkinElmer). Mean fluorescent intensity was measured within a cuboid region of interest, located within the mesencephalon, identified as the region within the head of the larvae between the visually identifiable structures of the eye and otic capsule.

#### Seizure-like locomotion assay

Following dechorionation and 24 hr treatment with pregnenolone and/or 3MA, 3 dpf larvae were transferred to individual wells of a 96 square well plate in 200 μL. The plate was mounted in a DanioVision (Noldus) tracking system at 25 °C with the chamber light on and movement of the larvae was tracked for 20 minutes. Movement parameters of individual larvae were quantified using EthoVision XT (Noldus) software. Zebrafish were assayed for seizure-like activity as in the drug screen.

### RNA Sequencing

Following dechorionation and 24 hr pregnenolone treatment from 2 dpf, 3 dpf larvae were anaesthetised with 0.016 % tricaine (made in aquarium water, buffered with 4 % 1M Tris pH 9 and adjusted to pH 7 with NaOH) and transferred to a 50 mL petri dish in a drop of water for dissection to separate heads and tails, and remove the yolk. The larvae were dissected with a 0.13 - 0.17 mm thick square glass coverslip. Two transverse cuts were made: 1, caudal to the hindbrain and rostral to the heart; 2, caudal to the yolk sac. Heads and tails were separated and all water was removed before flash freezing in liquid nitrogen. Six replicates of each condition were treated and processed in batches on the same day within a two-hour period.

Samples were disrupted and lysed using the FastPrep-24 speed 6 2 x 60” in 500 μL of kit lysis buffer. RNA was extracted from 400 μL of lysate and eluted in 50 μL. The RNA was then quantified using Quant-it RiboGreen assay (Life Technologies) and underwent quality control on a Fragment Analyzer.

Libraries were prepared with 150 ng of RNA input using the QuantSeq 3’ mRNA-Seq Library Prep Kit (FWD) HT for Illumina (Lexogen) and then quantified with Quant-it Picogreen (Life Technologies). Library sizes were controlled with the High Sensitivity NHS Fragment Analysis kit on a Fragment Analyzer (Agilent). RNA sequencing was performed on HiSeq 2500 with Rapid V2 chemistry SR 65 cycles (Illumina) loaded with 3 % Phix, using two Flow cells.

Raw count table was produced by HTseq-count^21^ and differential expression analysis was performed using Bioconductor R package edgeR (version 3.36).^22^ Gene set enrichment analysis was performed using CAMERA (limma Bioconductor R package, version 3.50.3) with custom gene sets (Supplementary Tables 2 and 3).^23^

### Hormone analysis in zebrafish larvae

Dechorionated 48 hpf zebrafish were treated in groups of 25 with DMSO or pregnenolone for 24 hr. At 3 dpf, following treatment, two groups of samples were pooled so that 50 larvae were transferred to 1.5 mL cryotubes and all water was removed before flash freezing in liquid nitrogen. Samples were shipped on dry ice and further sample processing and hormone analysis using a Sciex QTRAP 6500+ mass spectrometry system was performed by The Metabolomics Innovation Centre, Canada. Hormone measurements were normalised to an average weight of 3 dpf WT or *tpp1^-/-^*mutant larvae.

### Statistical analysis

Data are expressed as box-and-whisker plot (minimum to maximum) or mean ± SD, as indicated. Statistical analysis for all data were performed using unpaired t-test or linear mixed model in GraphPad Prism 9 and SPSS respectively. Differences between groups were considered statistically significant when *P* < 0.05. Statistical significance is indicated in figures.

## Results

### Drug screen in *tpp1*^-/-^ zebrafish identifies pregnenolone as a hit compound reducing seizure-like locomotion and cell death

As locomotion of zebrafish is easily and rapidly monitored, locomotion in *tpp1^-/-^* zebrafish, was used as the primary quantitative measure to screen 640 FDA-approved compounds. Locomotion assays were supplemented with other quantitative and qualitative assays (Fig 1A, Supplemental Table 1). Among the 640 screened compounds, 34 of these were identified as potential hits due to improvement in at least one assay and re-tested using a new batch of compound. Among these, pregnenolone (from screen 2) was identified as the only hit due to apparent improvement in retinal size, seizure-like locomotion at 72 hpf, locomotion at 96 hpf and survival in 96 hpf. No other compounds showed a reduction in any phenotypes when re-tested. However, after testing pregnenolone under blind conditions, only a reduction in duration of movement in *tpp1*^-/-^ zebrafish in the seizure-like locomotion assay remained statistically significant. Blind testing therefore validated the effect of pregnenolone on seizure-like locomotion, as demonstrated by a significant reduction in movement duration, distance and velocity in *tpp1*^-/-^ zebrafish (Fig 1 B,C).

The anti-seizure effect of pregnenolone on *tpp1*^-/-^ zebrafish was verified by electroencephalogram (EEG) analysis at 3 dpf after a 24hr treatment with 4 µg/ml pregnenolone (Fig 1 D,E). Typical seizure-like brain activity in *tpp1*^-/-^ zebrafish can be seen in Fig 1D, showing single representative EEG recordings from *tpp1*^-/-^ zebrafish treated with DMSO or pregnenolone, alongside healthy WT siblings. Mean EEG amplitude from all measured *tpp1^-/-^* zebrafish (n=6 DMSO treated, n=8 pregnenolone treated) show a significant reduction in EEG amplitude in those treated with pregnenolone compared with the DMSO control. These results support the locomotion assay results described above, showing an anti-seizure effect of pregnenolone in *tpp1*^-/-^ zebrafish.

Although we did not detect a statistically significant improvement of other phenotypes in our re-screen and blind testing of pregnenolone, we wondered whether cell death in *tpp1^-/-^* zebrafish brains was altered by pregnenolone treatment. A TUNEL cell death assay revealed a significant reduction of cell death in the mesencephalon of *tpp1*^-/-^ zebrafish treated with pregnenolone for 3 hours from 52 hpf, compared with DMSO (Fig 1 F,G). This finding suggests pregnenolone results in an improvement in pathology as well as seizures.

### Pregnenolone ameliorates lysosomal structure and function in *tpp1*^-/-^ zebrafish

Pregnenolone is an endogenous neurosteroid, a direct metabolite of cholesterol and the precursor of all steroid hormones.^24,25^ Pregnenolone is also an agonist of the sigma-1 receptor, which binds and regulates cholesterol levels within subcellular compartments, especially in lipid rafts.^26^ Pregnenolone was recently identified in another screen for compounds improving lysosomal storage in CLN3 disease, also known as juvenile Neuronal Ceroid Lipofuscinosis: in ARPE-19 CLN3 knockout cells pregnenolone was shown to reduce storage accumulation in the lysosomes.^27^ We went on to investigate whether pregnenolone has a therapeutic effect on the cell pathology of *tpp1*^-/-^zebrafish. To this end, we generated transgenic zebrafish with ZsGreen-tagged lysosomal membrane protein Lamp1 under the muscle-specific *actc1b* promoter in the background of *tpp1*^-/-^ zebrafish. The muscle reporter provides a large area of homogenous tissue allowing for accurate high-throughput measurement of the number and average size of lysosomes in live zebrafish, which would be more complicated in brain tissue owing to its smaller size and vast differences in lysosomal function between different neuronal cell populations. Quantification of confocal images show a strong lysosomal phenotype in the muscle of *tpp1*^-/-^ zebrafish; the average size of lysosomes is significantly increased and the number of lysosomes is significantly decreased compared to healthy WT siblings (Fig 2 A,B). Both these parameters were significantly improved in *tpp1*^-/-^ zebrafish treated with pregnenolone for 24 hr, compared to the DMSO control. Interestingly, pregnenolone had the same effect on the lysosomes of WT zebrafish, namely a significant decrease in average lysosome size and significant increase in lysosome number (Fig 2 A,B).

**Figure 2:**
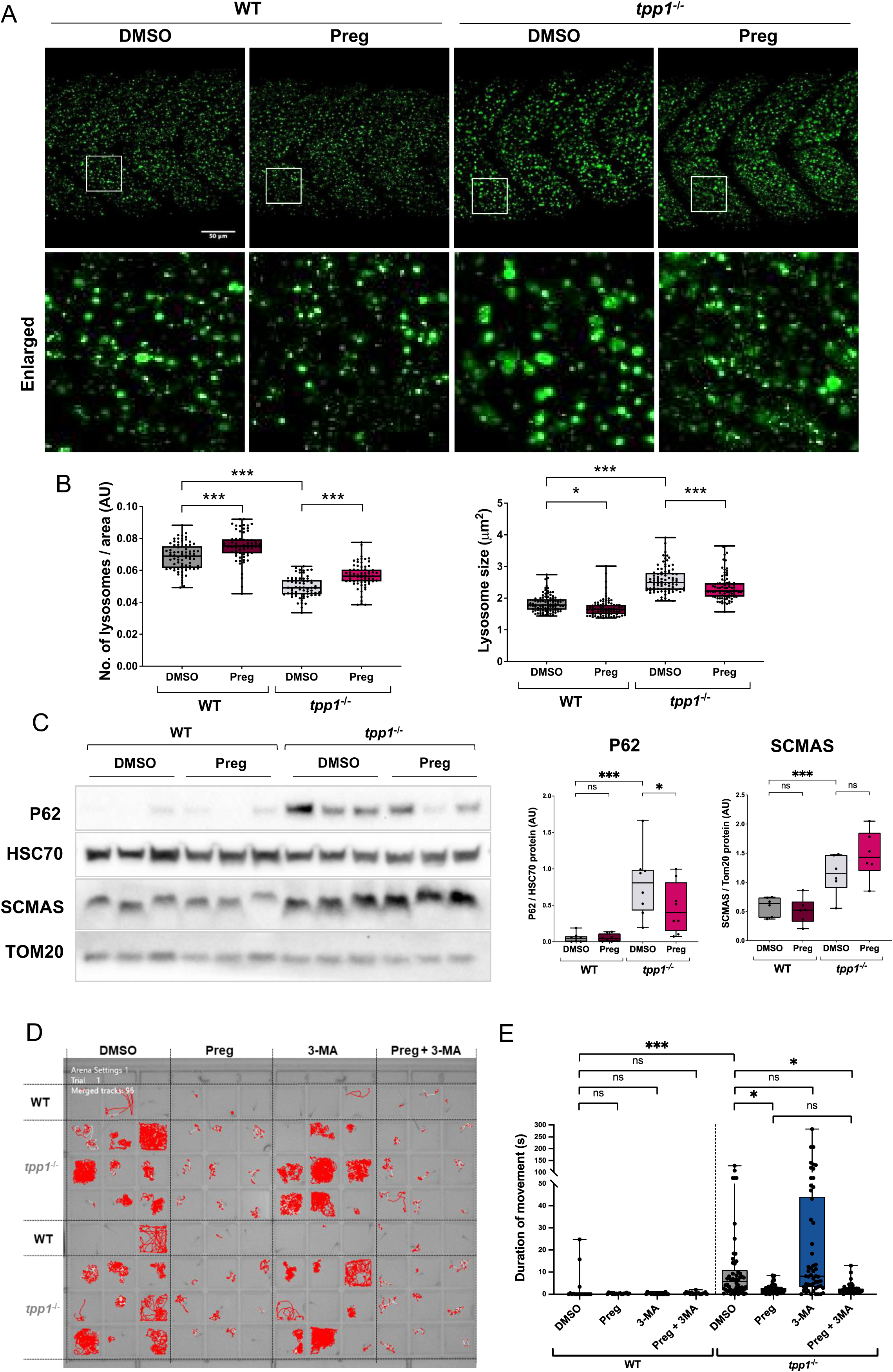
Pregnenolone ameliorates lysosomal abnormalities and dysfunction in *tpp1^-^*^/-^ zebrafish. (A-B) *ZsGreen-Lamp1* WT and *tpp1^-^*^/-^ zebrafish at 3 dpf treated with DMSO or pregnenolone (preg) for 24 hr. (A) Representative 40x max-projection confocal images, enlarged area indicated by white box. (B) Quantification of lysosomal number (left) and size (right) from ZsGreen-Lamp1 confocal images. Results from three replicate experiments. Data are shown by box and whisker plot (minimum to maximum) with individual points each representing an individual fish. (C) Representative image of Western blot analysis from whole lysates of de-yolked non-transgenic WT and *tpp1^-^*^/-^ zebrafish at 4 dpf treated with DMSO or preg for 48 hr, stained with antibodies against P62 normalized to HSC70 and SCMAS normalized to Tom20. Data are shown by box and whisker plot (minimum to maximum) with individual points each representing an individual sample. (D,E) Seizure-like locomotion assay from 3 dpf WT and *tpp1^-^*^/-^ mutant zebrafish treated with pregnenolone (Preg) + 3-MA for 24 hr, with DMSO, pregnenolone, or 3-MA alone as controls. (D) Representative locomotion assay plate. 96-well plate containing one fish per well in 200 µL aquarium water with treatment. Locomotion tracks are shown in red. White tracks are predicted movements that were not measured. (E) Plot showing duration of movement (s). No significant difference between pregnenolone with and without 3-MA, in *tpp1^-/-^*zebrafish. Results from three replicate experiments. Data are shown by box and whisker plot (min to max) with individual points each representing an individual fish. (B, C) Linear mixed model (fixed: genotype, treatment / random: date of experiment/plate). (E) Linear mixed model on log transformed data (fixed: genotype, treatment / random: date of experiment). *P\** <0.05, *P\*\**<0.01, *P\*\*\** <0.001.

Autophagy is a key function of the lysosome and a process that has been shown to be impaired in various LSDs, including CLN2 disease.^28–32^ The neuroprotective effect of autophagy induction and the inverse detriment caused by defects in autophagy have been widely demonstrated in the literature among a multitude of neurodegenerative diseases.^30,33,34^ Given that we see a decrease in lysosomal size with pregnenolone treatment, we measured P62, an autophagic adaptor protein, to test whether pregnenolone treatment enhances clearance of autophagic material in the lysosome, leading to the observed reduction in lysosomal size. Western blot analysis of de-yolked zebrafish tissue revealed a significant accumulation of P62 in *tpp1*^-/-^ zebrafish compared to WT siblings at 4 dpf (treated with DMSO), which was significantly reduced by 48 hr pregnenolone treatment (Fig 2C). Western blot analysis showed pregnenolone had no effect on the accumulation of SCMAS, the major component of storage material in CLN2 disease (Fig 5A). RNA sequencing analysis of *p62* shows that pregnenolone does not alter mRNA quantity at 3 dpf following 24 hr treatment, suggesting the reduction in P62 protein accumulation is indicative of enhanced clearance in *tpp1*^-/-^ zebrafish (Supplementary Fig 1A). Similarly, mRNA quantity of three different SCMAS loci, *atp5mc1*, *atp5mc3a* and *atp5mc3b,* is also unaffected by pregnenolone treatment (Supplementary Fig 1B), indicating that the lack of change in SCMAS protein represents no change in clearance.

Reduction in the accumulation of P62 protein in *tpp1*^-/-^ zebrafish treated with pregnenolone compared with DMSO suggests enhanced autophagic clearance. To test this hypothesis further, we generated transgenic zebrafish with ZsGreen-tagged autophagosome marker LC3, again under the muscle-specific *actc1b* promoter in the background of *tpp1*^-/-^ zebrafish. Autophagic flux was measured in zebrafish by live imaging of ZsGreen-LC3 with and without a 4 hr treatment of NH_4_Cl to block lysosomal degradation. At 3 dpf, *tpp1^-/-^* zebrafish did not yet show altered ZsGreen-LC3 flux compared with WT. However, preliminary results from one experiment showed that both WT and *tpp1*^-/-^ zebrafish revealed a significant accumulation of ZsGreen-LC3 following 24 hr pregnenolone treatment in the presence of NH_4_Cl, compared to the DMSO + NH_4_Cl control (Supplementary Fig 2 A,B). These results, in combination with the reduction of P62 protein accumulation, suggest pregnenolone treatment induces autophagy in both WT and *tpp1^-/-^*zebrafish.

To test whether the induction of autophagy by pregnenolone is responsible for the alleviation of seizures in *tpp1*^-/-^ zebrafish, we performed a locomotion assay on the zebrafish with pregnenolone treatment in combination with autophagy induction inhibitor, 3-MA.^35^ Results from 3 dpf zebrafish following 24 hr concurrent treatment of pregnenolone and 3-MA show duration of movement is comparable to pregnenolone treatment only (Fig 2 D,E). This finding therefore suggests that induction of autophagy is not the mechanism of action of pregnenolone’s anti-seizure effect.

### Pregnenolone acts via an mTORC1/TFEB-independent mechanism

Lysosomal biogenesis and function are largely regulated by mammalian target of rapamycin 1 (mTORC1) and transcription factor EB (TFEB), which respond to nutrient queues at the surface of the lysosome to regulate lysosomal gene expression and autophagy.^36,37^ Given the effect on lysosomes in both WT and *tpp1^-^*^/-^ zebrafish, we set out to test whether pregnenolone treatment acts via an mTORC1/TFEB-dependent mechanism. Western blot analysis of mTORC1 metabolite S6, revealed significantly increased phosphorylation of S6 in *tpp1^-^*^/-^ zebrafish compared with WT, however pregnenolone had no effect on this result (Fig 3A). Furthermore, measurement of nuclear translocation of ZsGreen-TFEB using 3 dpf *ZsGreen-TFEB;mCherry-nls* transgenic zebrafish, as has been published previously,^16^ suggested suppression of TFEB nuclear translocation (Fig 3 B,C). 24 hr treatment with pregnenolone showed no effect in either WT or *tpp1^-^*^/-^ zebrafish. These results highlight mTORC1/TFEB signalling abnormalities in *tpp1^-^*^/-^ zebrafish, however pregnenolone functions independently of this pathway. Enrichment analysis of a lysosomal gene set (Supplementary Table 2) supported these results, showing upregulation of lysosomal genes in *tpp1^-^*^/-^ zebrafish compared with WT, which was not further altered by pregnenolone treatment (Fig 3 D,E).

**Figure 3:**
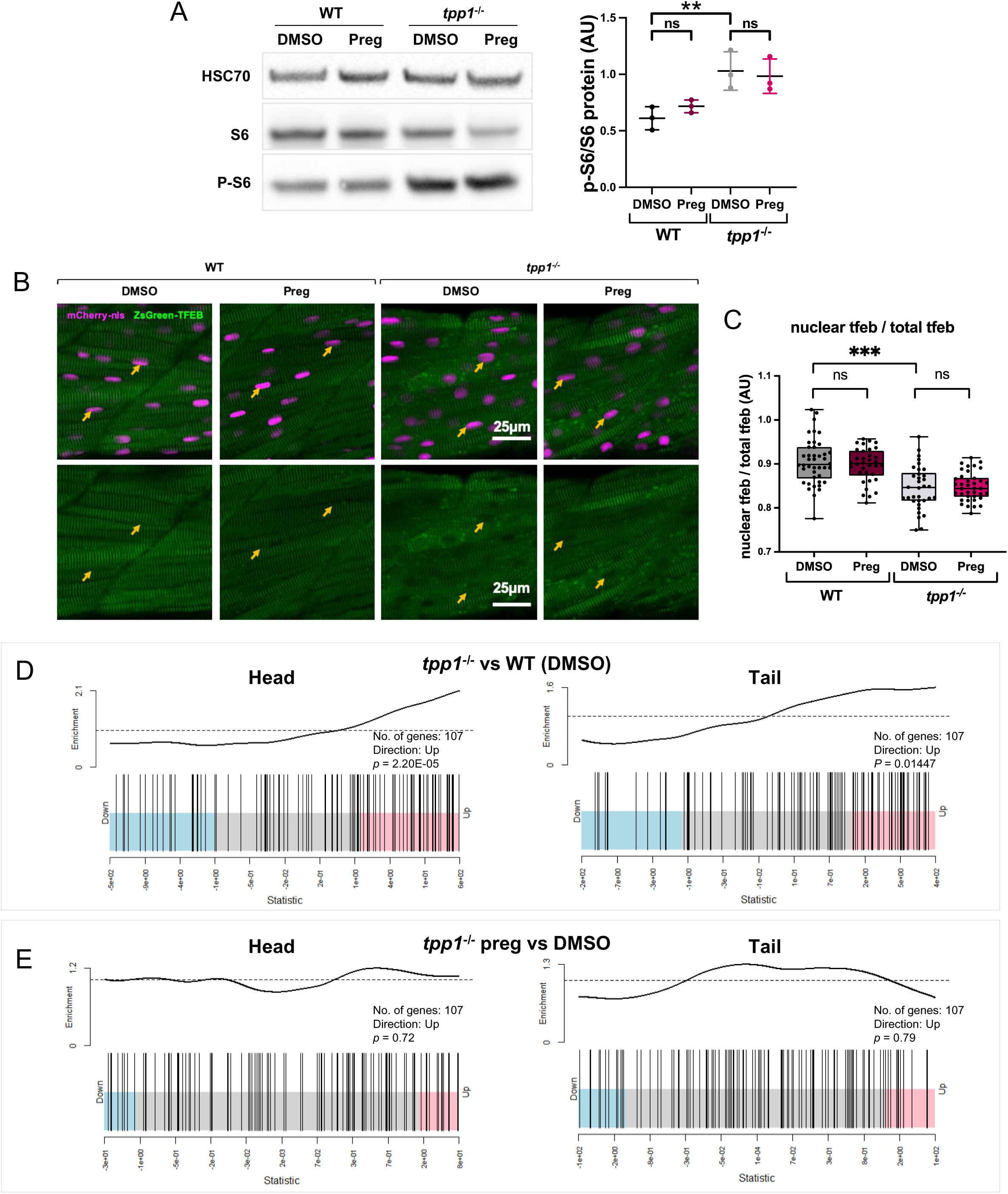
Lysosomal abnormalities and dysfunction in *tpp1^-^*^/-^ zebrafish are ameliorated by pregnenolone. (A) Representative image of Western blot analysis from whole lysates of de-yolked non-transgenic WT and *tpp1^-^*^/-^ zebrafish at 3 dpf treated with DMSO or pregnenolone (preg) for 24 hr, stained with antibodies against S6 and phosphorylated(p)-S6 normalized to HSC70. Data are shown as mean ± SD with individual points each representing an individual sample. (B,C) *ZsGreen-TFEB;mCherry-nls* WT and *tpp1^-^*^/-^ zebrafish at 3 dpf treated with DMSO or preg for 24 hr. (B) Representative cropped 40x single-plane images showing distribution of ZsGreen-TFEB, alongside mCherry-nls nuclear marker. ZsGreen-TFEB/mCherry-nls merge shown with corresponding ZsGreen-TFEB-only image below. Examples of nucleus indicated by yellow arrow. (C) Quantification of ZsGreen-TFEB intensity within nucleus over intensity of total area of muscle measured. (A,C) Linear mixed model (fixed: genotype, treatment / random: date of experiment/plate). *P\** <0.05, *P\*\**<0.01, *P\*\*\** <0.001. (D,E) RNA Sequencing data of a custom lysosomal gene set analysed in head (left) and tail (right) tissue of 3 dpf (D) *tpp1*^-/-^ vs WT zebrafish (24 hr DMSO treatment) and (E) *tpp1*^-/-^ zebrafish with preg vs DMSO treatment (treated for 24 hr). Results from 6 replicate samples, n=10 fish per sample. Barcode plots from CAMERA gene set enrichment analysis. Each bar represents a single gene; red and blue sections are representations of the portion of genes that most contribute to up- and down-regulated enrichment respectively. The curves show the enrichment score of the bars in the barcode plot; parts above and below the dashed line signify up- and down-regulated enrichment respectively. CAMERA *p* value result and highest enriched direction indicated on each plot.

### Perturbance in cholesterol and hormone metabolism in *tpp1*^-/-^ zebrafish

Pregnenolone is a direct metabolite of cholesterol; the conversion of cholesterol to pregnenolone precedes the production of all steroid hormones and is a rate-limiting step in steroidogenesis.^38^ In this regard, we analysed the endogenous concentrations of pregnenolone and a number of downstream steroid hormones in WT and *tpp1*^-/-^ zebrafish, and assessed the effect of pregnenolone treatment on these. Hormone analysis by mass spectrometry was conducted in 3 dpf zebrafish following 24 hr treatment with pregnenolone or DMSO control. Results showed dysregulation of steroid hormones in *tpp1*^-/-^ zebrafish compared with WT siblings in control conditions (Fig 4A and supplementary Fig 3), including significantly higher levels of pregnenolone (Fig 4B). Four out of the twenty-two measured hormones – 18-OH-corticosterone, cortisol, cortisone and 20α-DH-cortisol – were decreased in *tpp1*^-/-^ zebrafish compared with WT siblings under DMSO conditions and were rescued by pregnenolone treatment (Fig 4B and supplementary Fig 4). Seven steroids were significantly increased in *tpp1*^-/-^ zebrafish compared with WT siblings: and eleven were unchanged (Supplementary Fig 3). The changes in steroid hormone levels are summarised in Fig 5.

**Figure 4:**
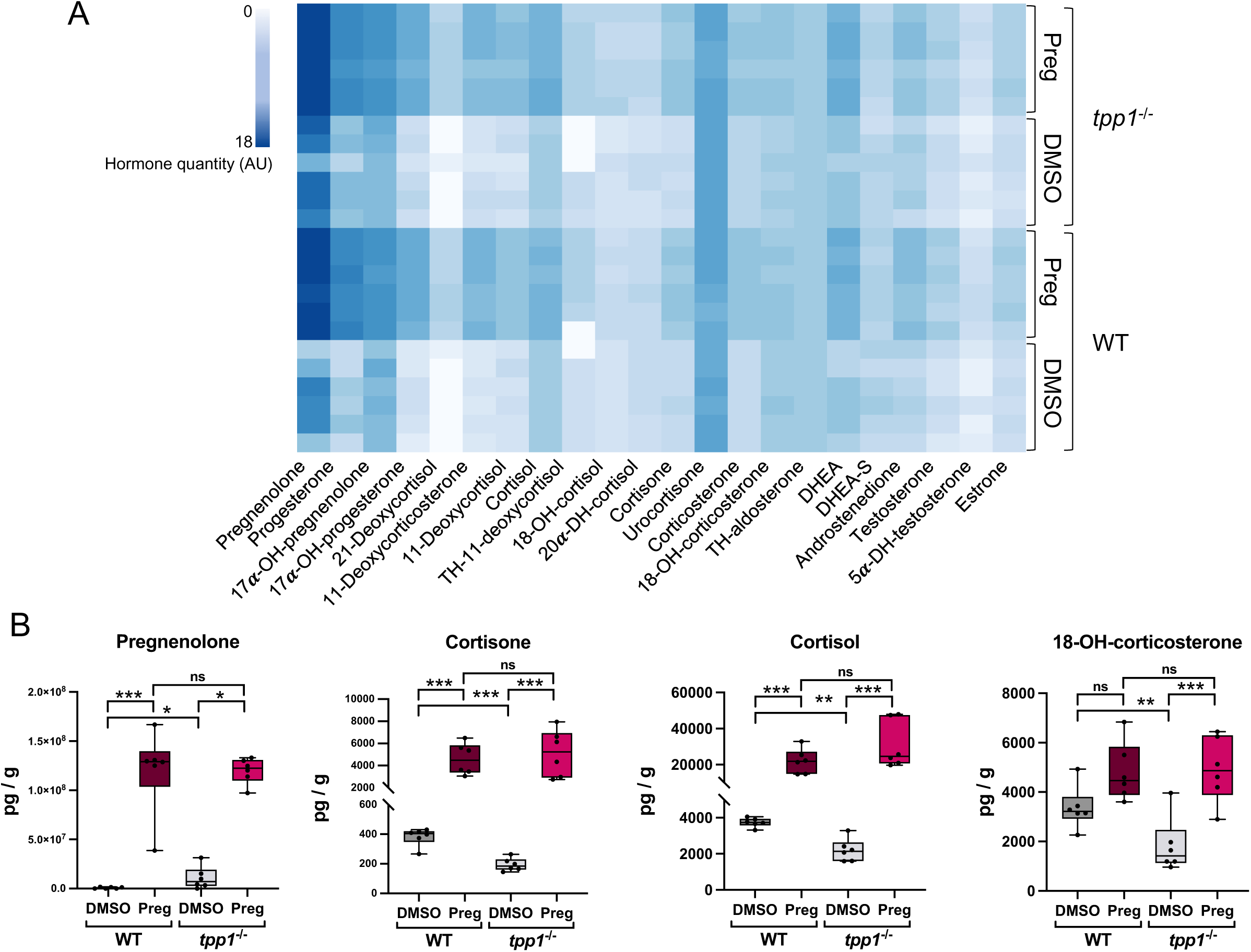
Steroid hormone biosynthesis is dysregulated in *tpp1*^-/-^ zebrafish and boosted by pregnenolone treatment. Steroid hormone measurements from 3 dpf WT and *tpp1*^-/-^ zebrafish treated with pregnenolone (preg) or DMSO for 24 hr. Results from 6 replicate samples, n=50 fish per sample. (A) Heatmap representing hormone measurements. Each block represents a single hormone from an individual sample. (B) Individual hormone measurement plots show picograms (pg) of hormones per gram of fish tissue. Data are shown by box and whisker plot (minimum to maximum) with individual points each representing an individual sample. Linear mixed model (fixed: genotype, treatment / random: date of experiment). *p\** <0.05, *p\*\**<0.01, *p\*\*\** <0.001.

**Figure 5:**
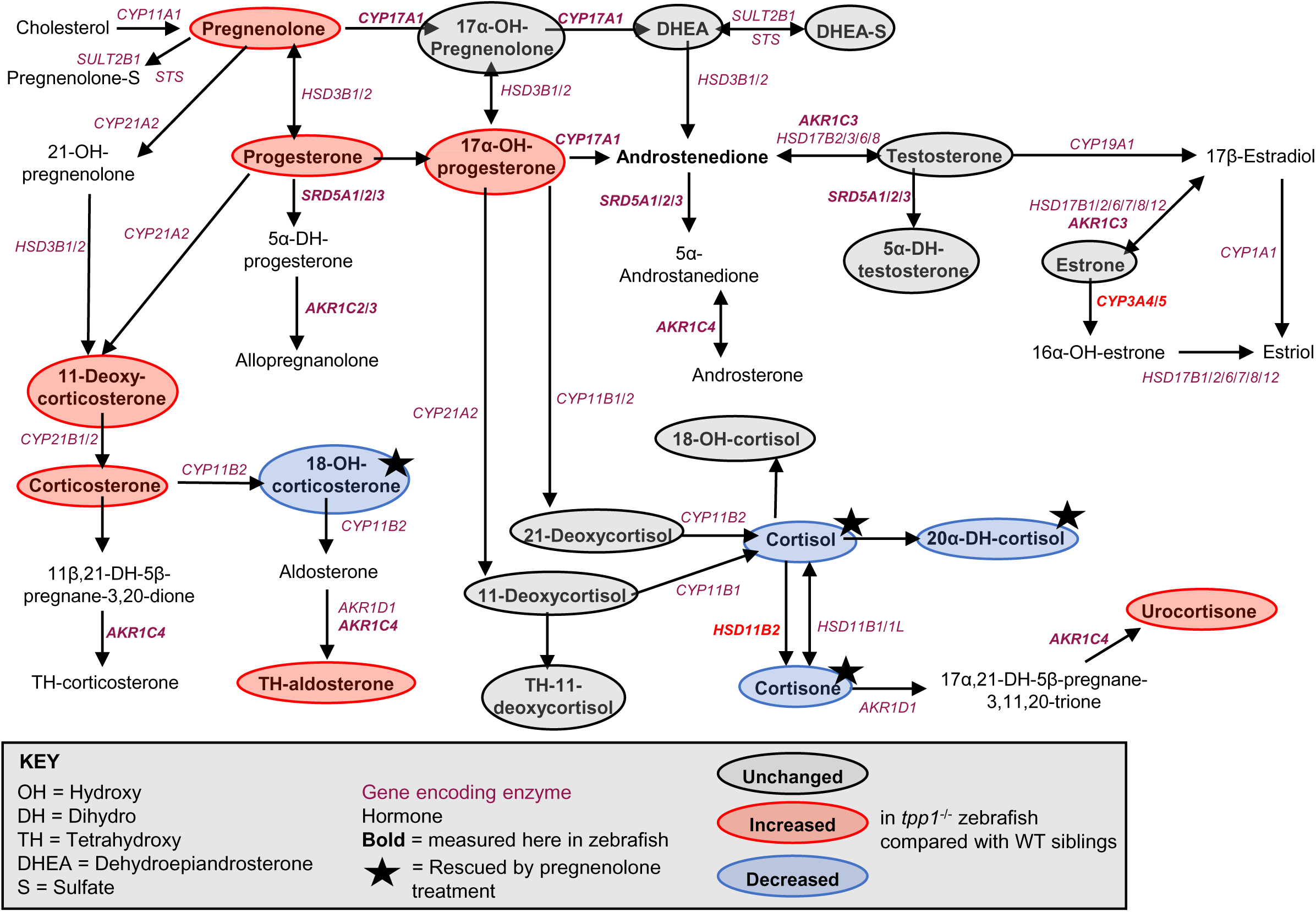
Overview of changes in steroid hormone levels in *tpp1^-^*^/-^ zebrafish compared with WT siblings. An overview of the steroid hormone synthesis pathway with blue and red circles representing hormones that were significantly increased or decreased respectively in *tpp1^-^*^/-^ zebrafish compared with WT siblings (treated with DMSO for 24 hr). Grey circles represent hormones that were unchanged. Hormones that are not circled were not measured. Hormones were measured by mass spectrometry in 3 dpf zebrafish (adapted from ^78^).

To further investigate the cholesterol and steroid pathway we performed enrichment analysis of a custom set of genes related to this pathway (Supplementary Table 3). The RNA Seq data from the zebrafish revealed significant enrichment of the cholesterol and steroid pathway in *tpp1*^-/-^ zebrafish compared with WT siblings in both head and tail tissue (Fig 6A). Pregnenolone treatment stimulated this pathway in WT zebrafish (Fig 6B) and lead to further enrichment of the pathway in *tpp1*^-/-^ zebrafish (Fig 6C).

**Figure 6:**
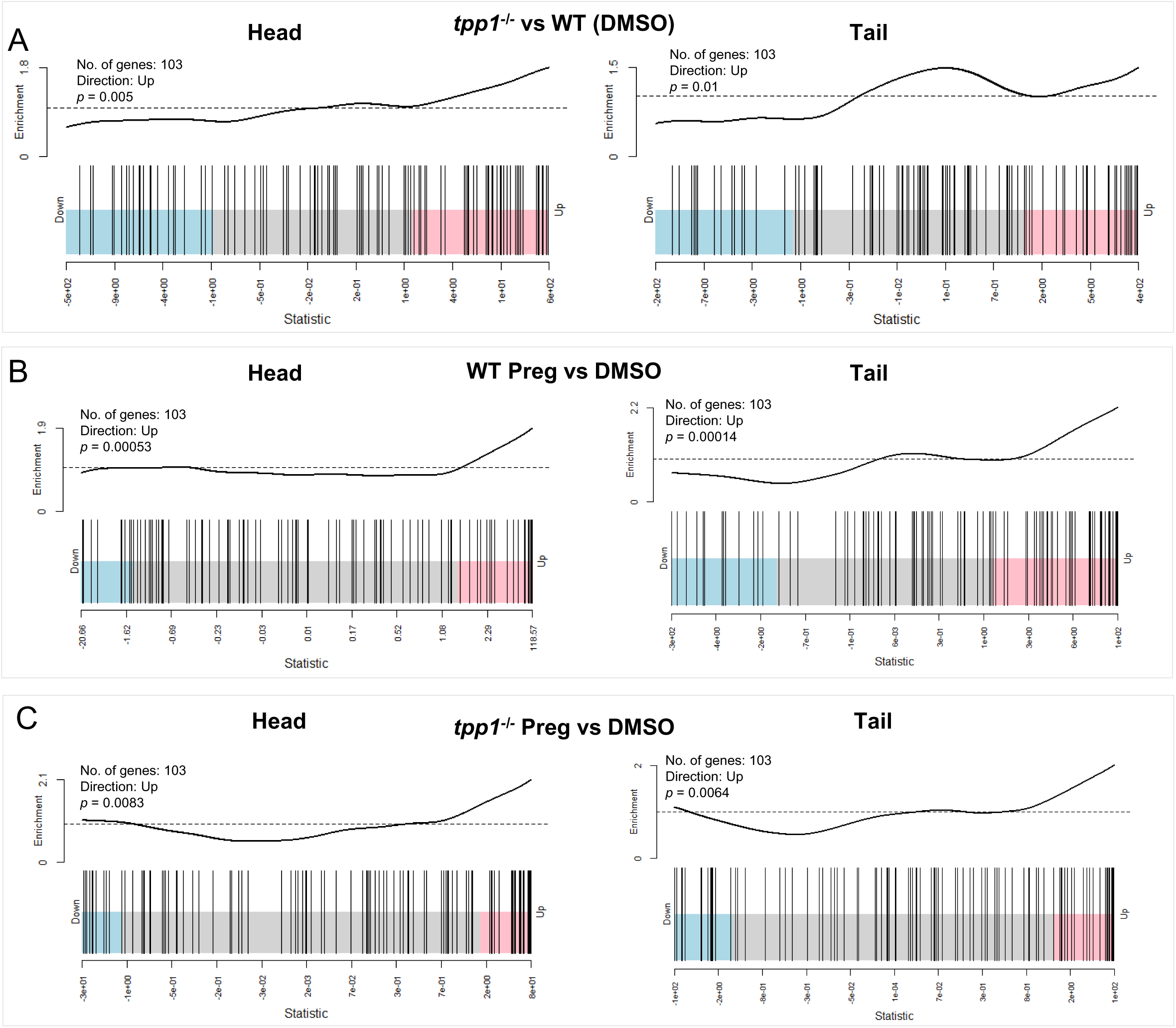
Cholesterol and hormone related genes are enriched in *tpp1*^-/-^ zebrafish and further boosted by pregnenolone treatment. RNA Sequencing data from head (left) and tail (right) tissue of 3 dpf zebrafish treated with DMSO or pregnenolone (Preg) for 24 hr. Results from 6 replicate samples, n=10 fish per sample. (A-C) Barcode plots from CAMERA gene set enrichment analysis. Each bar represents a single gene; red and blue sections are representations of the portion of genes that most contribute to up- and down-regulated enrichment respectively. The curves show the enrichment score of the bars in the barcode plot; parts above and below the dashed line signify up- and down-regulated enrichment respectively. CAMERA *p* value result and highest enriched direction indicated on each plot. (A) *tpp1*^-/-^ vs WT zebrafish (B) pregnenolone vs DMSO treated WT zebrafish (C) pregnenolone vs DMSO *tpp1*^-/-^ zebrafish.

Looking at individual genes involved in the various steps of cholesterol metabolism and hormone biosynthesis,^39^ *tpp1*^-/-^ zebrafish show significant changes in several major genes with diverse roles (Supplementary Fig 4, Supplementary Tables 4 and 5). These changes are summarised in Fig 7, which highlights the diverse functions within the cholesterol metabolism pathway of the proteins encoded by the mRNAs with altered expression in *tpp1*^-/-^ zebrafish and *tpp1*^-/-^ zebrafish treated with pregnenolone. Three genes are of particular interest because their mRNA levels are increased in *tpp1*^-/-^ but brought nearer to normal levels by treatment with pregnenolone. These are *lcat*, *stard3* and *npc2*, encoding Lecithin-cholesterol acyltransferase (LCAT), StAR-Related Lipid Transfer Domain Containing 3 (StARD3) and Niemann-Pick C2 (NPC2). StARD3 resides on the late endosome/lysosome membrane and NPC2 is an internal late endosome/lysosome protein, but both are both important for the transfer of cholesterol from late endosomes/lysosomes to other subcellular membrane compartments such as the plasma membrane, ER and mitochondria.^40,41^ LCAT, however, is found in the ER where it converts free cholesterol to cholesterol esters for storage in lipid droplets.^42^

**Figure 7:**
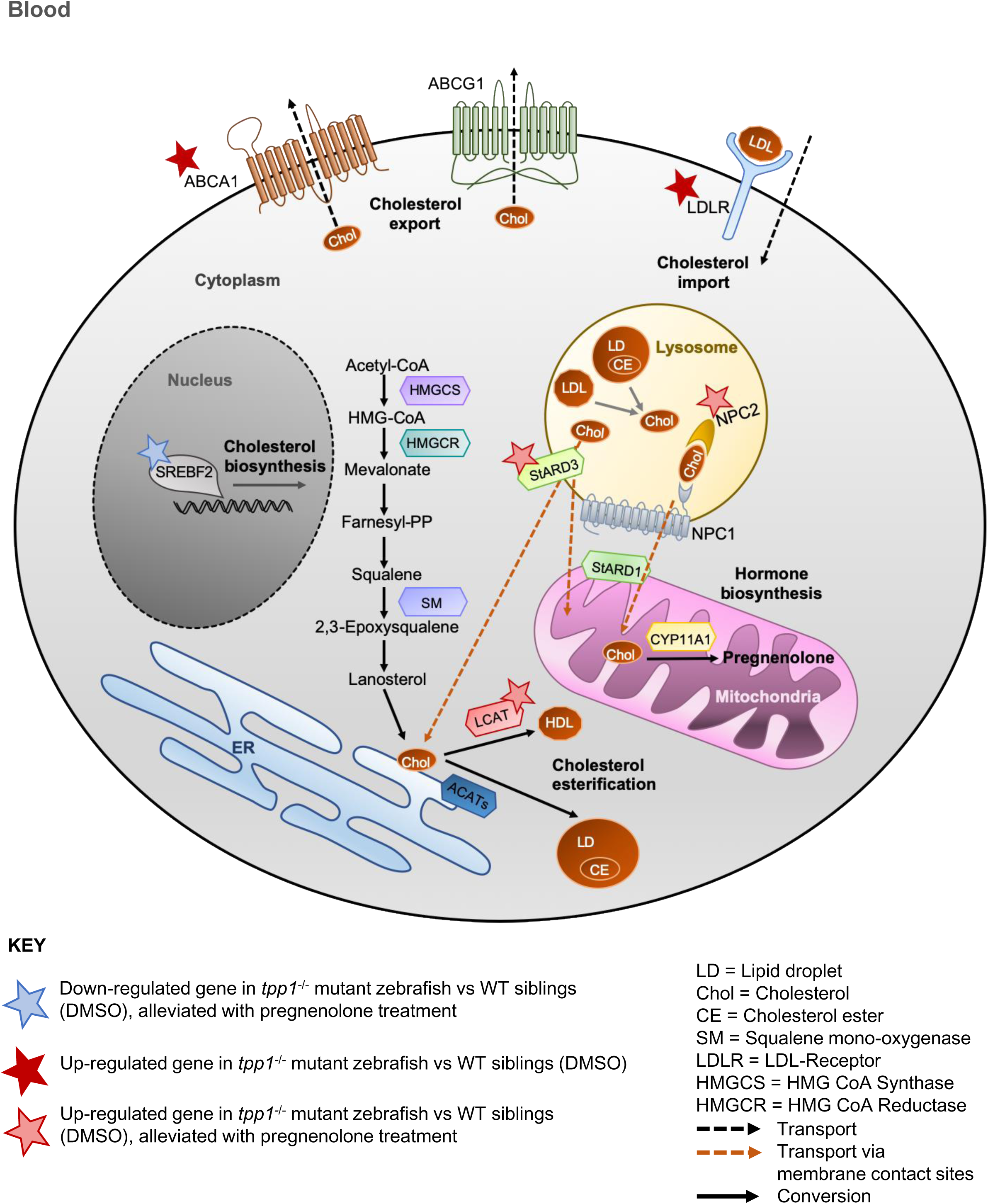
Overview of major pathways of cholesterol metabolism in the cell. Cholesterol availability in the cell depends predominantly on uptake of circulating LDL by the LDL-Receptor (LDLR) and de novo biosynthesis from acetyl-coA, mediated by the rate-limiting enzymes 3-hydroxy-3-methylglutaryl coenzyme A reductase (HMGCR) and squalene mono-oxygenase (SM). Cholesterol biosynthesis is regulated by the transcription factor SREBF2. Excess cholesterol is exported to the blood through ABCA1 or ABCG1. Imported or stored cholesterol esters are unesterified in the lysosome and free cholesterol can be distributed to other organelles in the cell via membrane contact sites, mediated by NPC1 and NPC2, and StARD3. Free cholesterol can be delivered to the inner mitochondrial membrane, mediated by StARD1, for use in hormone biosynthesis. Excess cholesterol can be esterified by ACAT or LCAT for storage in lipid droplets or excretion as HDL (adapted from^39,40,79,80^).

## Discussion

Seizures are a major symptom of CLN2 disease that worsen with disease progression and present a challenge for management of the condition as they often become resistant to anti-epileptic drugs.^9,10^ The present study demonstrates the use of the seizure-like locomotion assay in the *tpp1*^-/-^ zebrafish model of CLN2 disease to identify potential candidate compounds to alleviate seizures. Our screen identified pregnenolone as a hit compound, shown here to elicit an anti-seizure effect in *tpp1*^-/-^ zebrafish. Furthermore, reduced cell death in the CNS of *tpp1*^-/-^ zebrafish treated with pregnenolone reveal a neuroprotective effect. The results here show a partial rescue of lysosomal impairment in *tpp1*^-/-^ zebrafish treated with pregnenolone. This is evident from improved morphology of the lysosome in muscle cells of the zebrafish and a decrease in P62 accumulation following treatment of pregnenolone. These results present pregnenolone as a powerful candidate for further testing as a novel therapeutic for CLN2 disease. Pregnenolone is an FDA-approved compound and has been shown to be safely administered in adults for other conditions e.g.,^43,44^ and is now being trialled in children 14 years or older,^45^ thus pregnenolone could be considered for orphan designation for CLN2 disease.^46^

In line with our results, pregnenolone was recently identified in another screen, in ARPE-19 CLN3 knockout cells, where it was shown to reduce storage accumulation in the lysosomes.^27^ Our preliminary LC3 flux results suggest storage reduction is associated with induction of autophagy by pregnenolone. Although we do not see a reduction in SCMAS protein with pregnenolone treatment in the *tpp1*^-/-^ zebrafish, the decrease in lysosomal size and reduction of P62 protein accumulation suggest an overall reduction in storage accumulation. SCMAS is believed to be a direct substrate of TPP1/Tpp1^4,47–49^ and Tpp1 is almost entirely absent in *tpp1*^-/-^ zebrafish^14^; thus, the persistent accumulation of SCMAS despite improvements in lysosomal function is unsurprising. This observation raises the question of whether autophagy induction, by pregnenolone or other means, would provide a long-term therapeutic effect, or if the lysosomes would quickly be overwhelmed with SCMAS accumulation, despite a general improved clearance of other storage material.

Autophagy induction has been shown to have significant therapeutic effects in other models of LSDs caused by specific enzyme deficiency. In a *Drosophila* model of Gaucher disease lacking the glucocerebrosidase enzyme, for example, induction of autophagy by means of mTORC1 inhibition by rapamycin treatment was shown to significantly increase lifespan and improve locomotor and oxidative stress phenotypes.^50^ Extended lifespan and phenotypic improvement by autophagy induction in the Gaucher disease model indicates a sustained therapeutic effect that can lead to significant improvements in disease outcome. Despite these benefits associated with autophagy induction, inhibiting autophagy with 3-MA in *tpp1*^-/-^ zebrafish here did not affect pregnenolone’s anti-seizure effect, indicating that autophagy is unlikely to be the mechanism of action responsible for this particular therapeutic effect. Further confirmation is needed to ensure that 3-MA blocks pregnenolone-induced autophagy, although the treatment used clearly blocks autophagy induction in zebrafish.^18^

Pregnenolone might act via other mechanisms to induce the anti-seizure effect. Potential mechanisms include modulation of GABA_A_ receptors by downstream metabolites of pregnenolone^51,52^ and/or agonistic effects on the sigma-1 receptor, which has been shown to elicit neuroprotective effects.^53–55^ Neurosteroids are known for their role in modulation of neuronal excitability which has important implications in seizure susceptibility and control. Allopregnanolone, progesterone, androsterone and 11-deoxycorticosterone are all known to enhance GABA_A_ receptor function, eliciting anti-convulsant actions.^52,56^ Several studies have demonstrated the anti-seizure effect of allopregnanolone, and Ganaxolone, an analogue of allopregnanolone, has shown promising results in clinical trials for a number of epilepsies.^51,52^ These findings present an alternative mechanism for the seizure rescue observed in *tpp1*^-/-^ zebrafish treated with pregnenolone, as raised pregnenolone levels might lead to increases in anti-convulsant hormones. Consistent with this hypothesis, we see an increase in the concentration of a number of hormones following pregnenolone treatment, including anti-convulsant steroids, progesterone and 11-deoxycorticosterone. GABA_A_ activation has also been shown to have neuroprotective effects,^57–59^ thus, action through GABA_A_ signaling could also be responsible for the attenuation of cell death in *tpp1*^-/-^ zebrafish treated with pregnenolone.

Pregnenolone and its downstream metabolite DHEA, as well as their sulfated forms, act as sigma-1 receptor agonists.^54,55^ The sigma-1 receptor is an anti-epileptic target. Positive modulators of the receptor, fenfluramine and E1R, have been shown to elicit anti-convulsant effects.^60,61^ Fenfluramine has demonstrated substantial efficacy in reducing seizures in a phase 3 clinical trial for patients with Dravet syndrome, a refractory paediatric epilepsy.^62^ This finding suggests that actions through the sigma-1 receptor could therefore underlie the anti-seizure effect in *tpp1*^-/-^ zebrafish treated with pregnenolone.

The sigma-1 receptor is widely expressed in different tissues, including brain and muscle. It binds cholesterol and regulates a variety of protein binding partners in a variety of membranes.^26,63^ Sigma-1 receptor localisation to mitochondrial-associated membrane (MAM) microdomains of the ER is most studied. Here it regulates calcium flux into the mitochondria via the inositol trisphosphate receptor 3 (InsP_3_R3) calcium channel.^26,64^ Calcium regulation is involved in various cell death pathways and sigma-1 agonists have been shown to attenuate neuronal death in a number of models of neurodegenerative diseases.^26,65^ In a cell model of Huntington’s disease for example, the sigma-1 receptor agonist PRE-084 (2-(4-morpholinoethyl)-1-phenylcyclohexane1-carboxylate hydrochloride) increased antioxidants and reduced reactive oxygen species (ROS).^66^ In line with a role in regulating oxidative stress, knockdown of the sigma-1 receptor has been shown to increase ROS production, likely through dysregulation of calcium homeostasis.^67–69^ As increased levels of ROS have been documented in CLN2 disease patient cells,^32,70^ and oxidative stress can lead to cell death,^71^ pregnenolone might ameliorate cell death in *tpp1*^-/-^ zebrafish by alleviating oxidative stress through its action on the sigma-1 receptor. At the plasma membrane, the sigma-1 receptor decreases voltage-gated sodium and calcium channel activity whilst increasing activity of some voltage-gated potassium channels, thereby limiting excitotoxicity.^26^ This relationship provides an additional possible mechanism of action of pregnenolone in limiting both seizures and cell death in *tpp1^-/-^* zebrafish.

Identification of pregnenolone as a hit compound led us to investigate the cholesterol and steroid hormone pathway in our zebrafish model because 1) pregnenolone is a steroid that is synthesised from cholesterol in the mitochondria, and 2) cholesterol is bound by the sigma-1 receptor, for which pregnenolone is an agonist. This investigation led to findings of dysregulation throughout the cholesterol and steroid hormone pathways that have not been previously noted in CLN2 disease. We found increased endogenous pregnenolone in *tpp1^-/-^* zebrafish. As pregnenolone reduces seizures in those zebrafish, pregnenolone is the precursor of many steroids, and pregnenolone and those steroids can have anti-epileptic effects (see earlier), this increase in endogenous pregnenolone could be a compensatory mechanism. On the other hand, given that pregnenolone is a metabolite of cholesterol, elevated levels of endogenous pregnenolone in *tpp1^-/-^*zebrafish could result in depleted cholesterol stores, and pregnenolone treatment may reduce cholesterol depletion in addition to negating the need for the proposed compensatory increase in pregnenolone. In *tpp1^-/-^* zebrafish, we also see upregulation of *npc2*, *stard3* and *lcat* mRNAs and their restoration to normal mRNA levels by pregnenolone treatment. The effect on cholesterol storage and sub-cellular levels of free cholesterol resulting from these changes is difficult to predict, so further experiments will be needed. However, it is very likely that cholesterol handling is affected. Cholesterol is crucial for membrane organisation, neuronal differentiation and myelination.^72,73^ Furthermore, cholesterol is important for the integrity of the lysosomal membrane and modulates ion permeability.^74^ Disruption to the cholesterol pathway could contribute to CLN2 disease pathology via disturbance of its function in any one of these roles, highlighting cholesterol abnormality as an important area for further research in *tpp1^-/^*^-^ zebrafish and other models of CLN2 disease.

In summary, the present study employed a zebrafish model of CLN2 disease to perform a medium-throughput drug screen which identified pregnenolone as a potential therapeutic for the disease. Pregnenolone was shown to reduce seizures in our model and improve cell pathology, including improvement in lysosome morphology and function and reduced cell death. The safety profile of pregnenolone make it an appealing candidate for further testing into the use of it as a therapeutic for CLN2 disease and perhaps other NCLs.^43,44^ The novel *Lamp1-ZsGreen* transgenic zebrafish presented here revealed a striking phenotype in *tpp1^-/-^*zebrafish compared to healthy WT siblings. This transgenic line provides a powerful tool to investigate potential therapeutics that can improve lysosomal biology *in vivo*, as demonstrated by the effect of pregnenolone. The use of the *act1b* muscle promoter provides a substantial area of tissue that can be automatically detected and imaged using a high-content confocal imaging system. These lines can therefore be used as the basis of a screen similar to the one presented in the current study, through automated imaging in a 96 well format. There are a number of examples of similar high-content screening studies using other transgenic zebrafish lines as effective platforms for drug screening.^75–77^

## Data availability statement

RNAseq and metabolomic datasets will be uploaded to. Other data will be provided by the corresponding author on request.

## Acknowledgements

We thank Andrew Hibbert at the Royal Veterinary College for confocal imaging assistance and Yu Mei (Ruby) Chang for advice on statistical analysis.

