## Supplementary figures and legends for "Whole-organism screening in a zebrafish model of CLN2 disease identifies pregnenolone as a modulator of lysosomal functions with anti-epileptic properties"

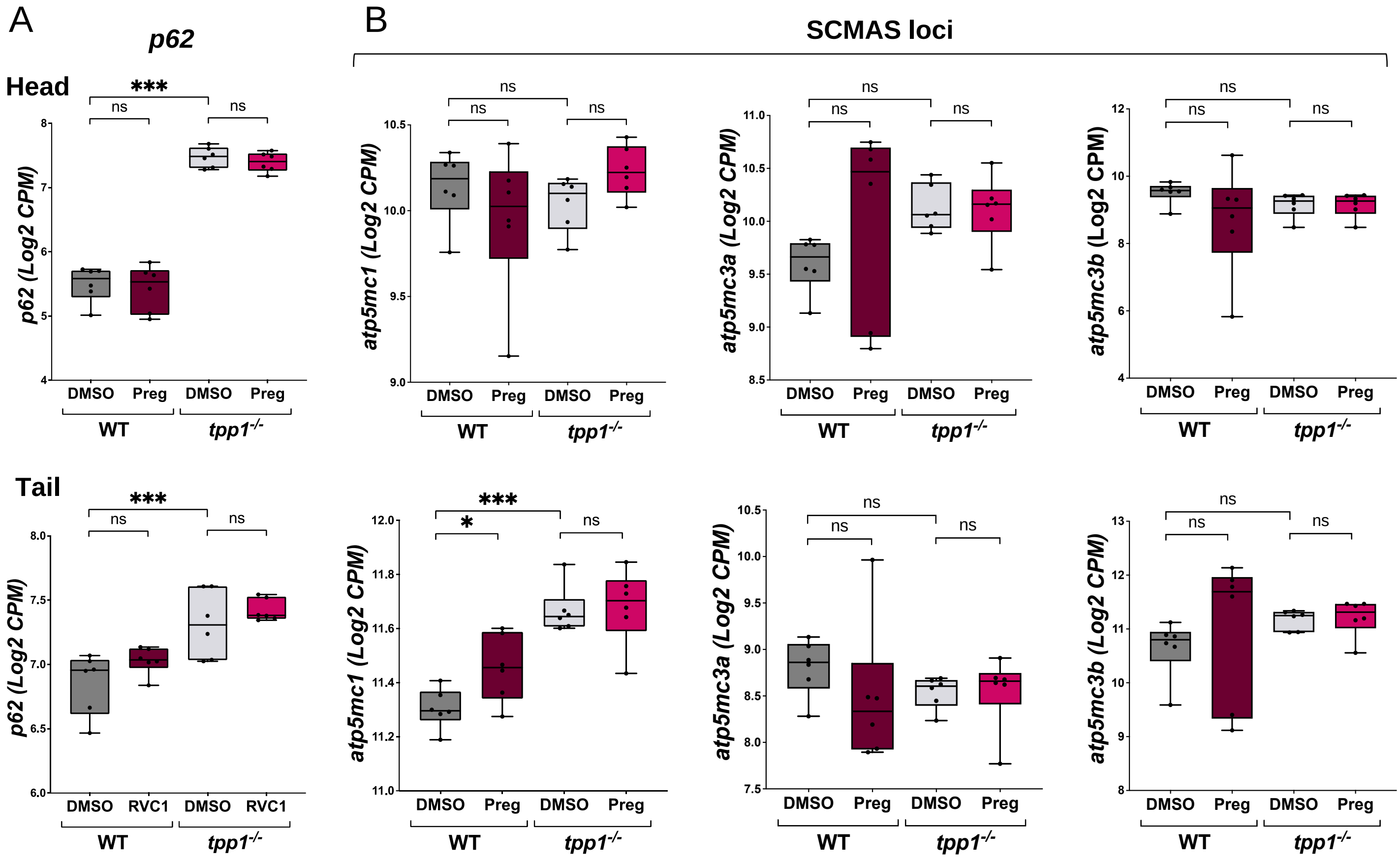


**Supplementary Figure 1: *p62* and SCMAS loci mRNA analysis.** RNA Sequencing data from head (top panel) and tail (bottom panel) tissue of 3 dpf WT and *tpp1*$^{-/-}$ mutant zebrafish treated with DMSO or pregnenolone (Preg) for 24 hr. Results from 6 replicate samples, n=10 fish per sample. Plots show Log2-transformed counts per million (CPM) for (A) *p62*; (B) SCMAS loci: *atp5mc1*, *atp5mc3a*, *atp5mc3b*. Data are shown by box and whisker plot (min to max) with individual points each representing an individual sample. Linear mixed model (fixed: genotype, treatment / random: plate). *P** <0.05, *P***<0.01, *P**** <0.001.

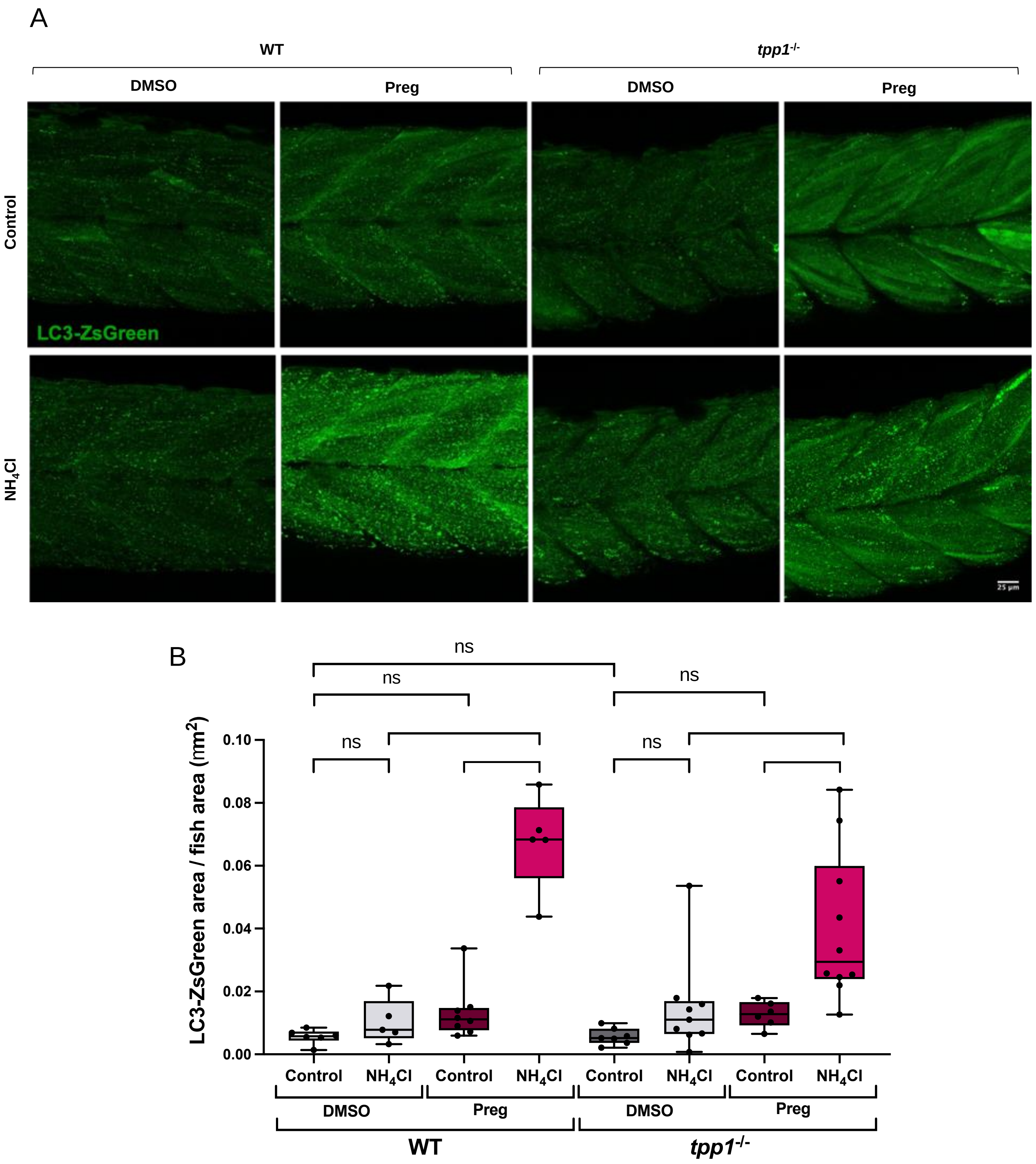


**Supplementary Figure 2: Preliminary ZsGreen-LC3 flux assay suggests pregnenolone increases autophagic flux in WT and *tpp1$^{-/-}$* mutant zebrafish.** *ZsGreen-LC3* WT and *tpp1$^{-/-}$* mutant zebrafish at 3 dpf treated with DMSO or pregnenolone (Preg) for 24 hr, with $NH_4Cl$ or $H_2O$ control for the last 4 hr. (A) Representative 40x max-projection images. (B) Quantification of ZsGreen-LC3 confocal images: increase in ZsGreen-LC3 area in both WT and *tpp1$^{-/-}$* mutant zebrafish treated with pregnenolone + $NH_4Cl$, compared to pregnenolone + control demonstrates increase in autophagic flux with pregnenolone treatment. Results from one experiment. Data are shown by box and whisker plot (minimum to maximum) with individual points each representing an individual fish. Linear mixed model (fixed: genotype, treatment / random: date of experiment). $p$* <0.05, $p$**<0.01, $p$*** <0.001.

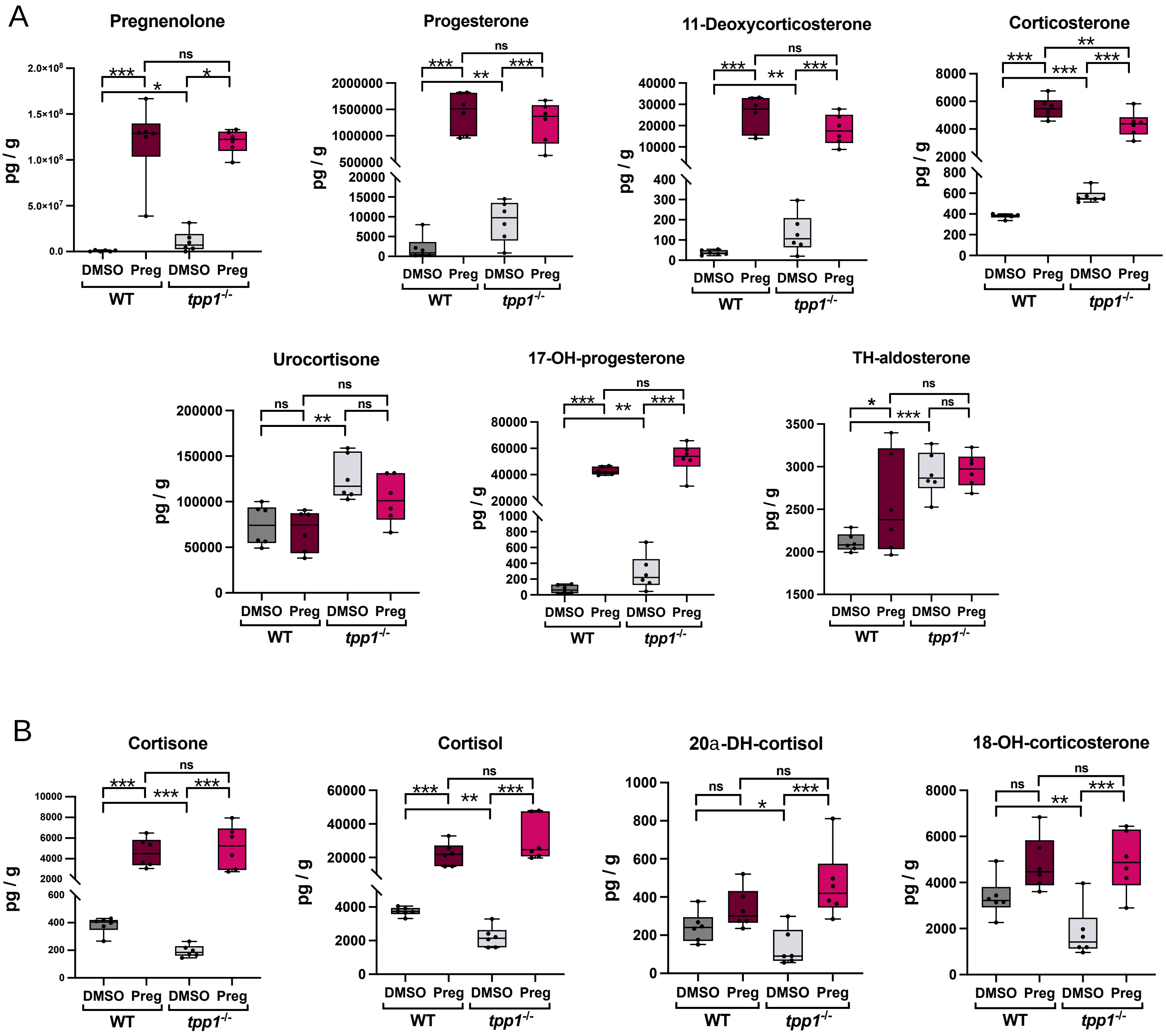
A
Pregnenolone
Progesterone
11-Deoxycorticosterone
Corticosterone
Urocortisone
17-OH-progesterone
TH-aldosterone
pg / g
DMSO Preg DMSO Preg
WT
tpp1-/-
ns
***
**
*
B
Cortisone
Cortisol
20a-DH-cortisol
18-OH-corticosterone

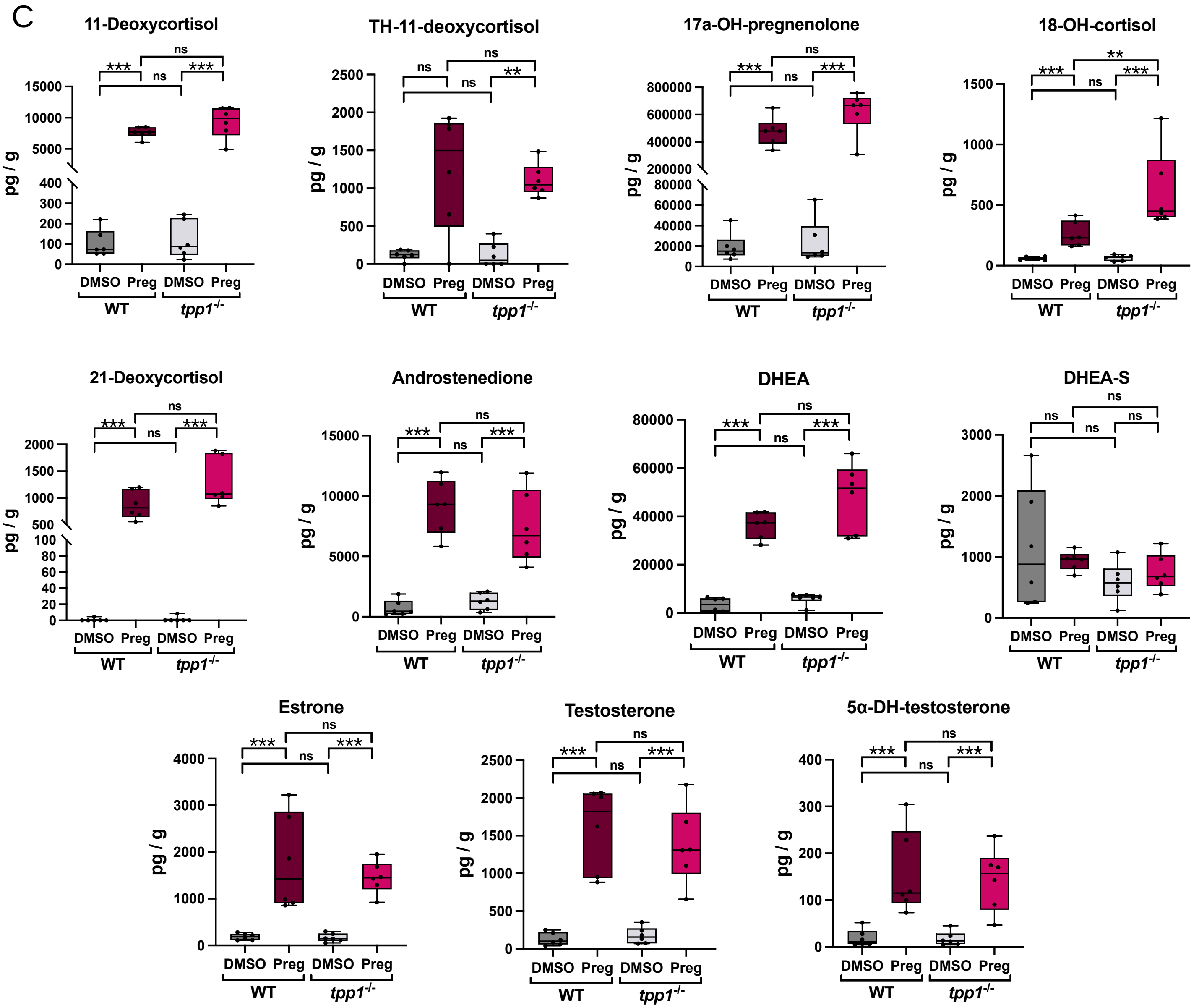


**Supplementary Figure 3 Hormone measurements in WT and *tpp1*$^{-/-}$ mutant zebrafish.** Hormone measurements from 3 dpf WT and *tpp1*$^{-/-}$ mutant zebrafish treated with pregnenolone (Preg) or DMSO for 24 hr. Results from 6 replicate samples, n=50 fish per sample. Individual hormone measurement plots show picograms (pg) of hormones per gram of fish tissue. Hormones are grouped by those that are increased (A), decreased (B) or unchanged (C) in *tpp1*$^{-/-}$ mutant zebrafish compared with WT (treated with DMSO). Data are shown by box and whisker plot (minimum to maximum) with individual points each representing an individual sample. Linear mixed model (fixed: genotype, treatment / random: date of experiment). *P** <0.05, *P***<0.01, *P**** <0.001.

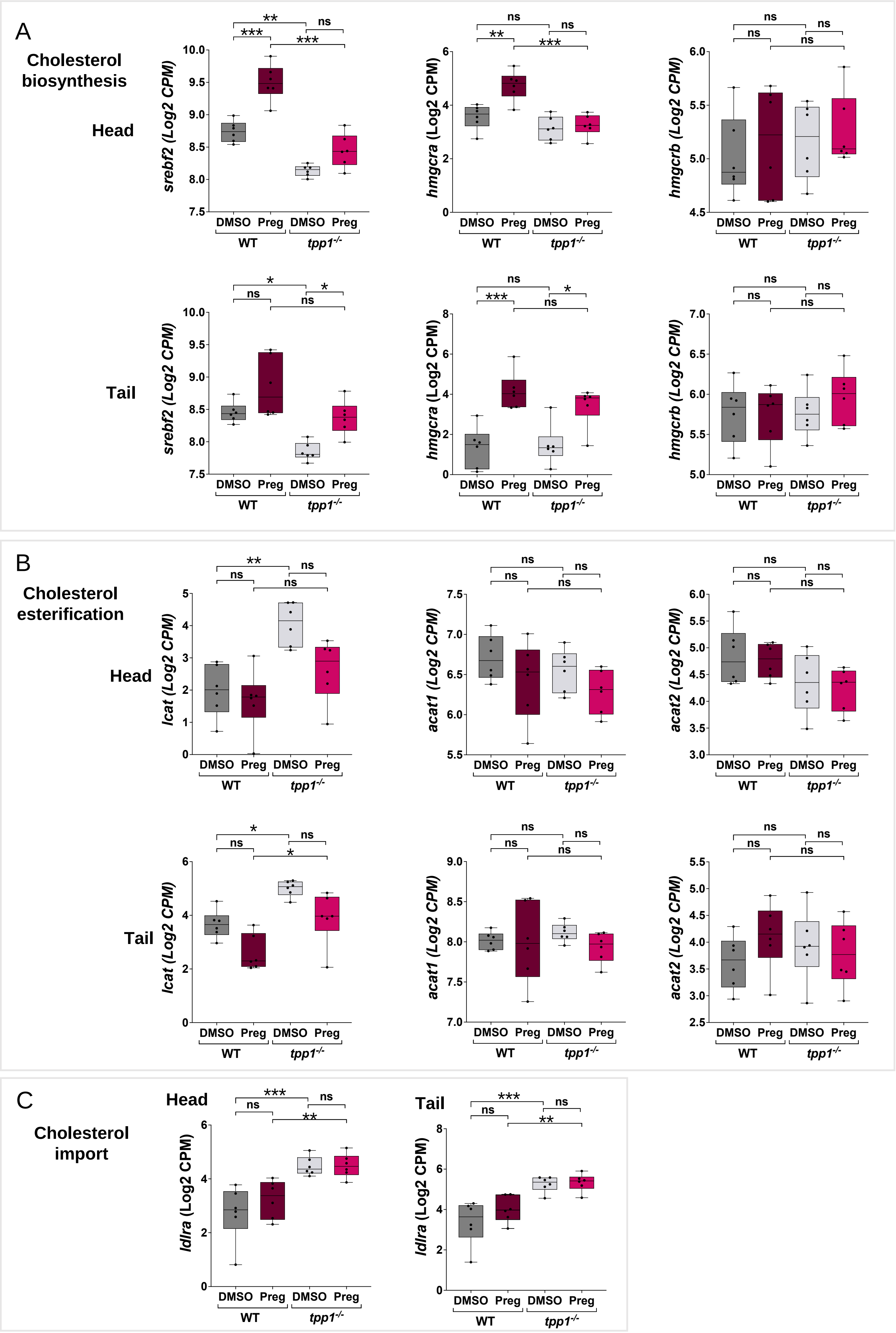

A
Cholesterol biosynthesis
Head
Tail
srebf2 (Log2 CPM)
hmgcra (Log2 CPM)
hmgcrb (Log2 CPM)
DMSO Preg DMSO Preg
WT
tpp1-/-
**
***
*
ns
B
Cholesterol esterification
Head
Tail
lcat (Log2 CPM)
acat1 (Log2 CPM)
acat2 (Log2 CPM)
C
Cholesterol import
Head
Tail
ldlra (Log2 CPM)

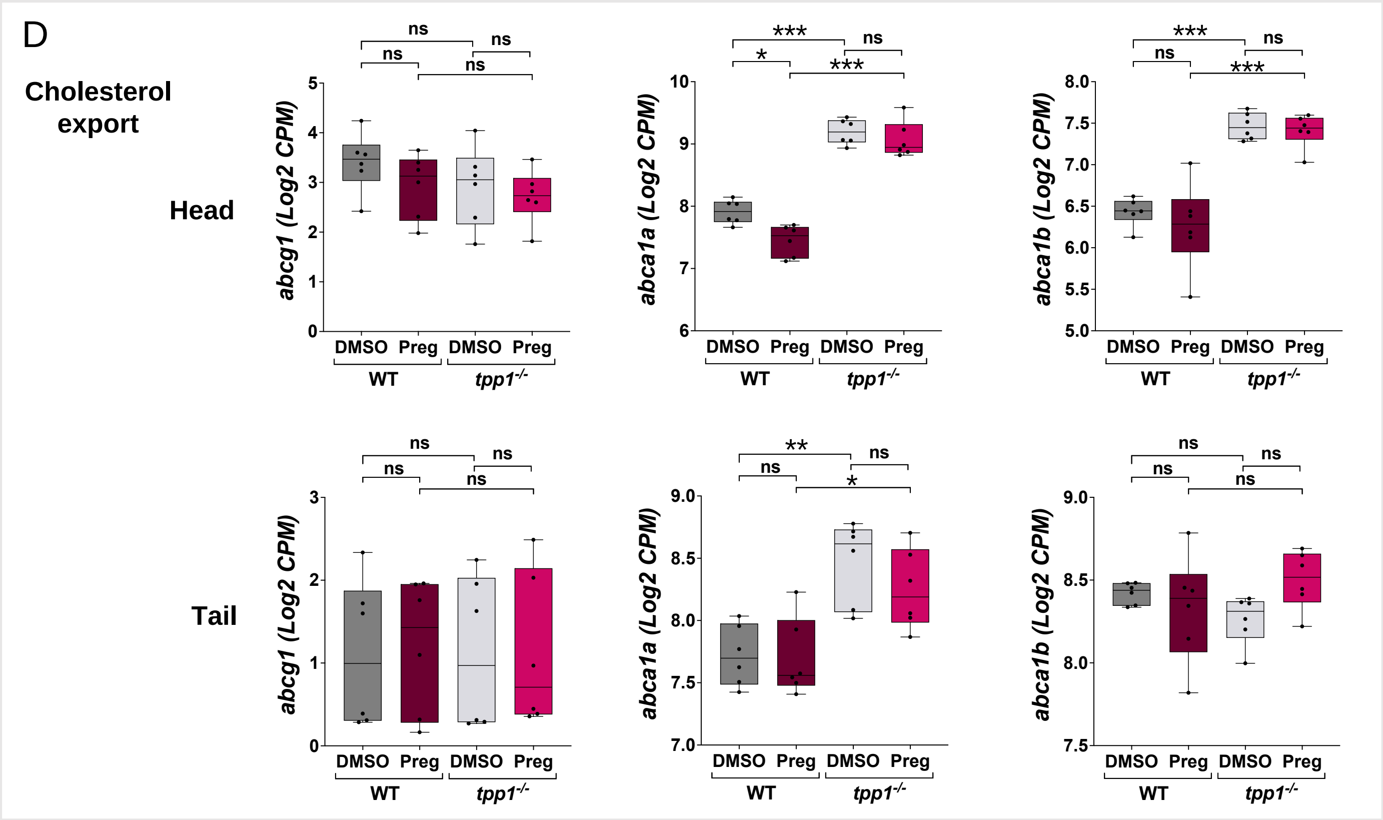
D
Cholesterol export
Head
Tail
abcg1 (Log2 CPM)
abca1a (Log2 CPM)
abca1b (Log2 CPM)
ns
*
**
***
DMSO
Preg
WT
tpp1-/-

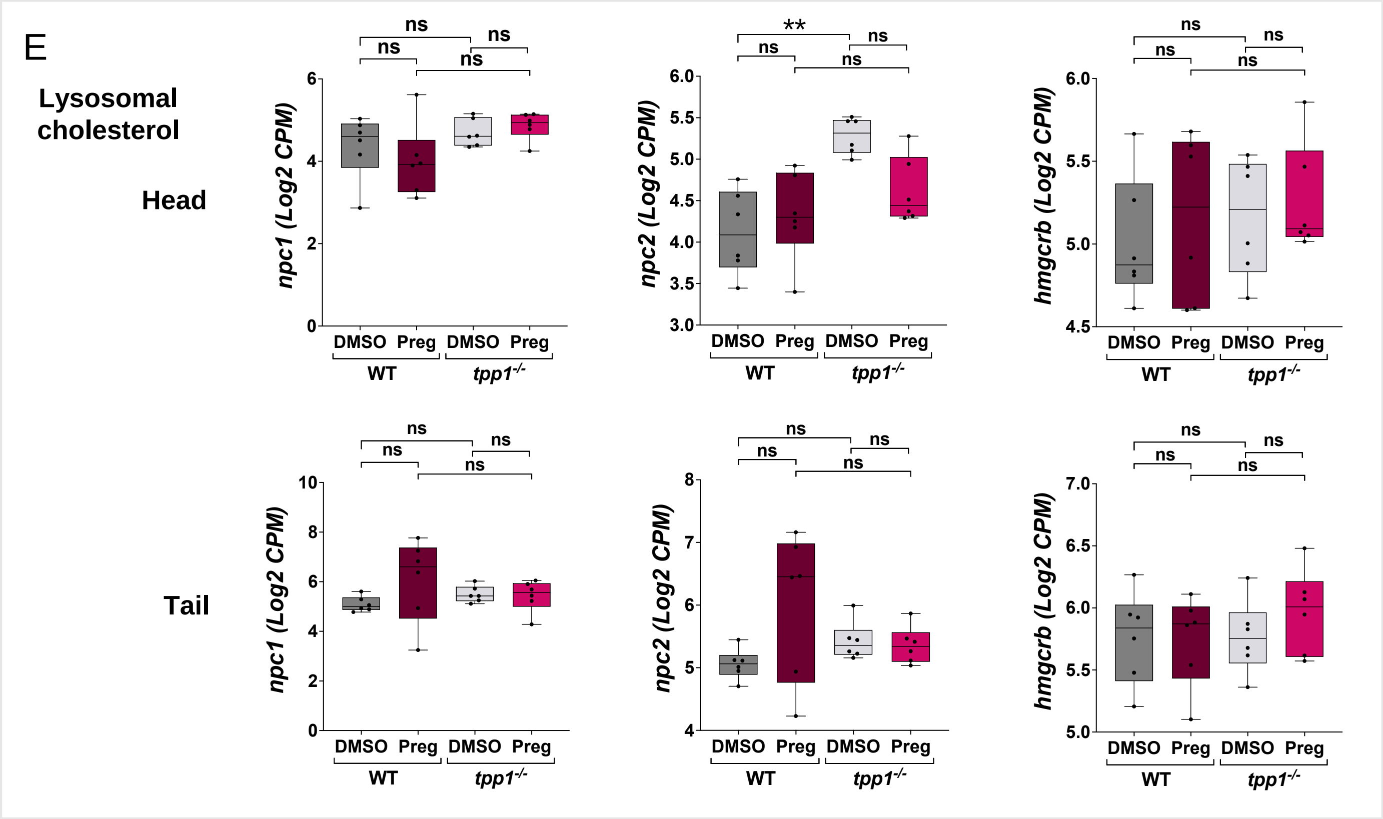
E
Lysosomal cholesterol
Head
Tail
npc1 (Log2 CPM)
npc2 (Log2 CPM)
hmgcrb (Log2 CPM)
ns
**
DMSO
Preg
WT
tpp1-/-

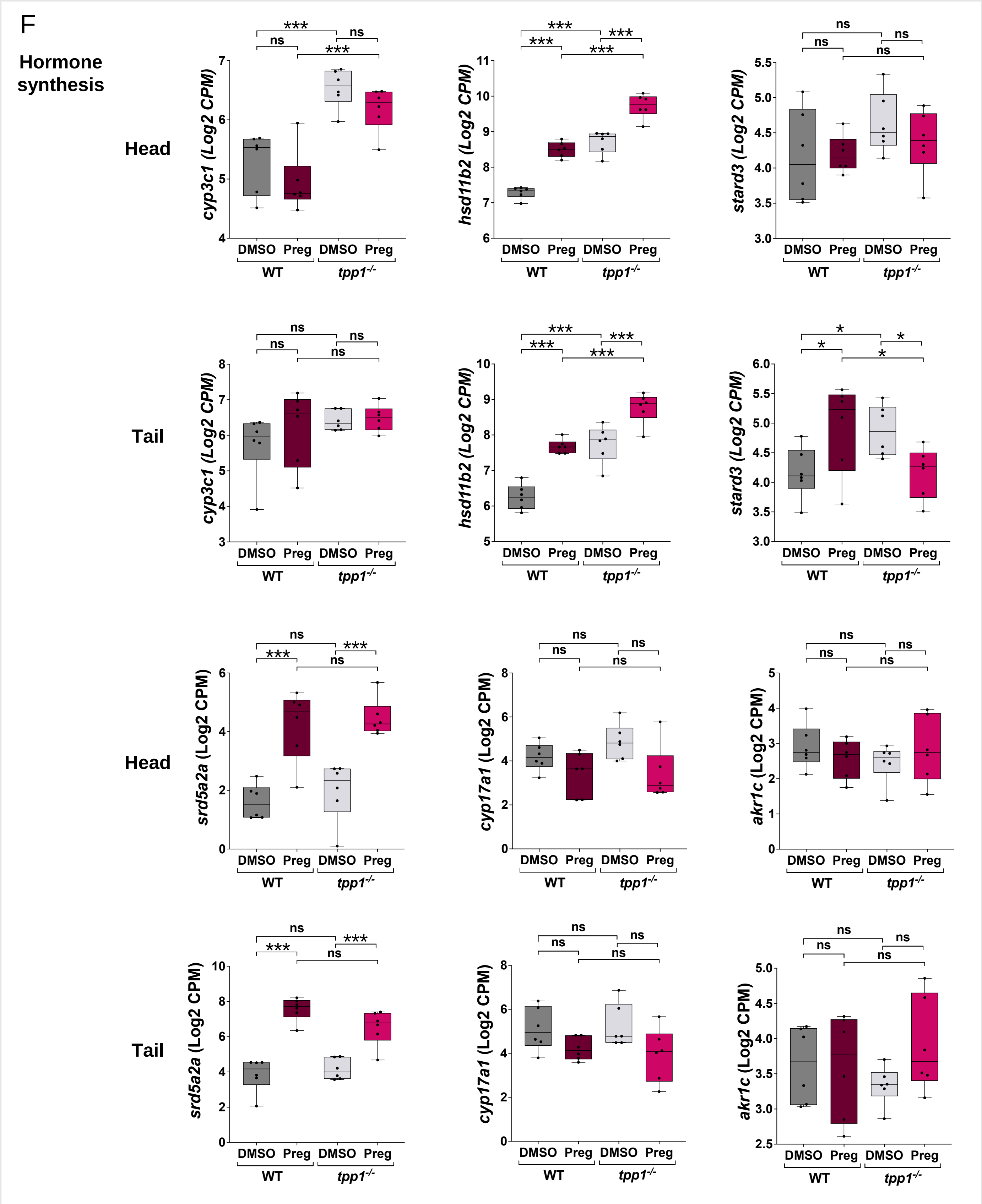


**Supplementary Figure 4: Dysregulation of key genes in the cholesterol and hormone pathway in *tpp1*$^{-/-}$ mutant zebrafish and the effect of pregnenolone (Preg) treatment.** (A-G) Individual gene analysis from RNA Sequencing data in head and tail tissue. Plots show Log2-transformed counts per million (CPM) for genes encoding key proteins involved in (A) cholesterol biosynthesis (*srebf2, hmgcra, hmgcrb*), (B) esterification (*lcat, acat1, acat2*), (C) import (*ldlra*) and (D) export (*abcg1, abca1a, abca1b*), (E) regulation of lysosomal cholesterol (*npc1, npc2*) and (F) hormone synthesis (*cyp3c1*, *hsd11b2*, *stard3*, *srd5a2a, cyp17a1, akr1c*), as indicated by left panel. Data are shown by box and whisker plot (minimum to maximum) with individual points each representing an individual sample. Linear mixed model (fixed: genotype, treatment / random: plate). $p^*$ <0.05, $p^{**}$<0.01, $p^{***}$ <0.001.
