## Supplementary tables for "Whole-organism screening in a zebrafish model of CLN2 disease identifies pregnenolone as a modulator of lysosomal functions with anti-epileptic properties"

Supplemental Table 1.

Two or three *tpp1-/-* zebrafish were treated with each drug from the FDA-approved drug library in one of two screens (Screen 1 and 2 respectively). Improvements in phenotypes were noted in several compounds from Screen 1 (A) and Screen 2 (B). Compounds in Tables A and B were re-sourced and re-tested using 10 *tpp1-/-* zebrafish but otherwise the same protocol as before. Only one compound from Screen 2 (pregnenolone) still resulted in phenotypic improvements (C). Performing the assays blinded in replicate using 10 *tpp1-/-* zebrafish per replicate revealed that pregnenolone only significantly improved seizure like locomotion at 76 hpf (D and Figure 1).

A. Summary of results from Screen 1

| Compound name | Retinal morphology at 120 hpf (qualitative) | Morphology at 120 hpf (qualitative) | Touch response at 120 hpf (quantitative) | Locomotion at 120 hpf (quantitative) | Survival at 120 hpf (quantitative) |
|---|---|---|---|---|---|
| Letrozole | | Improved | Improved | | |
| Cilnidipine | Improved | | | | |
| Pinacidil | | | | Improved | |
| Benserazide HCL | | | | Improved | |
| Spiperone | | | | Improved | |
| Flecainide | | | | Improved | |
| Acemetacin | | | | Improved | |
| Erlotinib | | | | Improved | |
| Cefepime | | | | Improved | |
| Didanosine | | | | Improved | |
| Eprosartan | | | | Improved | |
| Propofol | | | | Improved | |
| Oxaliplatin | | | | Improved | |
| Zafirlukast | | | | Improved | |
| Famotidine | | | | Improved | |

B. Summary of results from Screen 2 results.

| Compound name | Retinal size and morphology 76-96 hpf (qualitative) | Seizure-like locomotion at 76 hpf (quantitative) | Locomotion at 96 hpf (quantitative) | Survival at 96 hpf (quantitative) |
|---|---|---|---|---|
| Aceclofenac | Improved | Improved | Improved | Improved |
| Alendronate sodium | Improved | Improved | Improved | Improved |
| 2’,3’-dideoxycytidine | | Improved | Improved | Improved |
| Enrofloxacin | Improved | Improved | Improved | Improved |
| Lisinopril | Improved | Improved | Improved | Improved |

| | | | | |
|---|---|---|---|---|
| Triamcinolone | | Improved | Improved | Improved |
| Quinapril HCL | | Improved | Improved | Improved |
| Racecadotril | | | Improved | Improved |
| Ramipril | | Improved | Improved | Improved |
| Rimantadine HCL | | | Improved | Improved |
| Streptomycin sulphate | | Improved | Improved | Improved |
| Sulfadiazine | | | Improved | Improved |
| Sulfadimethoxine | | | Improved | Improved |
| Sulfasalazine | | Improved | Improved | Improved |
| Levofloxacin HCL | Improved | | Improved | Improved |
| Omeprazole | | | Improved | Improved |
| Oxiconazole nitrate | Improved | | Improved | Improved |
| **Pregnenolone** | **Improved** | **Improved** | **Improved** | **Improved** |
| Primaquine phosphate | | Improved | Improved | Improved |

C. Summary of results from re-testing compounds that cause phenotypic improvement in Screens 1 and 2.

| Compound name | Retinal size and morphology 76-96 hpf (qualitative) | Seizure-like locomotion at 76 hpf (quantitative) | Locomotion at 96 hpf (quantitative) | Survival at 96 hpf (quantitative) |
|---|---|---|---|---|
| **Pregnenolone** | | **Improved** | **Improved** | **Improved** |

D. Summary of results from blind testing of pregnenolone

| Compound name | Retinal size and morphology 76-96 hpf (qualitative) | Seizure-like locomotion at 76 hpf (quantitative) | Locomotion at 96 hpf (quantitative) | Survival at 96 hpf (quantitative) |
|---|---|---|---|---|
| **Pregnenolone** | | **Improved** | | |

**Supplemental table 2: Lysosome gene set**

| Gene symbol | GENE ID |
|---|---|
| Abca2 | 100006090 |
| acp1 | 541489 |
| Acp2 | 503759 |
| acp6 | 558758 |
| Aga | 566517 |
| Ap1b1 | 562166 |
| Ap1g1 | 324466 |
| Ap1g2 | 100007877 |
| Ap1m1 | 100330411 |
| Ap1m2 | 403021 |
| Ap1s1 | 393279 |
| Ap1s2 | 327237 |
| ap1s3a | 550415 |
| ap1s3b | 447897 |
| ap3b1a | 563316 |
| Ap3d1 | 563359 |
| Ap3m1 | 266798 |
| Ap3m2 | 415244 |
| Ap3s2 | 436812 |
| Ap4b1 | 393309 |
| Ap4s1 | 368852 |
| arsj | 559800 |
| asah1b | 393549 |
| atp6ap1a | 100004566 |
| atp6ap1b | 195824 |
| atp6ap1la | 100000415 |
| atp6v0a1a | 324307 |
| atp6v0a1b | 553691 |
| Atp6v0a2 | 561469 |
| atp6v0a2a | 415147 |
| Atp6v0b | 321724 |
| atp6v0ca | 192336 |
| atp6v0cb | 325402 |
| Atp6v0d1 | 322811 |
| Atp6v1h | 286779 |
| Cd164 | 555639 |

| | |
|---|---|
| Cln3 | 492340 |
| Clta | 406318 |
| Cltb | 100329879 |
| cltca | 323579 |
| cltcb | 503600 |
| cts12 | 567046 |
| cts12 | 567046 |
| ctsba | 406645 |
| Ctsc | 368704 |
| ctsd | 65225 |
| Ctsf | 565588 |
| Ctsh | 324818 |
| Ctsk | 550475 |
| ctsl.1 | 436641 |
| ctsla | 321453 |
| Ctsz | 450022 |
| dnase2 | 100000399 |
| Entpd4 | 436692 |
| fuca1.1 | 335494 |
| fuca1.2 | 445165 |
| Galns | 791159 |
| Gba | 559072 |
| Gga1 | 798361 |
| gga3b | 100334661 |
| Glb1 | 100001628 |
| Gm2a | 550529 |
| Gnptab | 553365 |
| gnsa | 566506 |
| gnsb | 327635 |
| Gusb | 571441 |
| Hexa | 550460 |
| Hexb | 323613 |
| Hgsnat | 100329505 |
| Ids | 559959 |
| Idua | 567720 |
| Igf2r | 557061 |
| lamp1b | 563328 |
| Lamp2 | 541406 |

| | |
|---|---|
| Laptm4a | 100003844 |
| Laptm4b | 368911 |
| Lgmn | 406625 |
| lipf | 406713 |
| Litaf | 431731 |
| M6pr | 406486 |
| Man2b1 | 541519 |
| Manba | 393128 |
| mcoln1a | 406689 |
| mcoln3a | 100334819 |
| Mfsd8 | 564342 |
| Naga | 560778 |
| Naglu | 560126 |
| Napsa | 336746 |
| Neu1 | 559850 |
| Npc1 | 553330 |
| Npc2 | 282673 |
| Pla2g15 | 335008 |
| PPID | 415155 |
| Ppt1 | 406648 |
| Psap | 140811 |
| scarb2a | 192340 |
| Sgsh | 563849 |
| si:ch211-212k18.7 | 100000085 |
| Slc11a2 | 678623 |
| smpd2b | 797914 |
| smpd3 | 568381 |
| smpd5 | 567592 |
| sort1a | 406511 |
| sort1b | 799153 |
| Sumf1 | 553423 |
| tcirg1a | 100003139 |
| tcirg1b | 406342 |
| Tpp1 | 798347 |

Supplemental Table 3: Cholesterol gene set

| Gene symbol | Gene ID |
|---|---|
| cyp17a1 | 399692 |
| sult2st3 | 777792 |
| sult2st2 | 777793 |
| srd5a2a | 550398 |
| cyp11c1 | 791124 |
| hsd11b1la | 393293 |
| hsd11b2 | 334098 |
| hsd17b7 | 768185 |
| hsd17b8 | 64815 |
| hsd17b12b | 322626 |
| hsd17b12a | 327417 |
| hsd20b2 | 368367 |
| zgc:92630 | 436969 |
| dhrs11a | 791578 |
| dhrs11b | 791770 |
| cyp3a65 | 553969 |
| cyp3c1 | 324340 |
| cyp1b1 | 100150054 |
| ugt2a4 | 553952 |
| ugt5e1 | 100002993 |
| ugt1b5 | 100384899 |
| comta | 561372 |
| cyp51 | 414331 |
| lbr | 368360 |
| ebp | 436600 |
| sc5d | 447891 |
| dhcr7 | 378446 |
| lipf | 406713 |
| cyp24a1 | 100004700 |
| abca1a | 558924 |
| abca1b | 100136868 |
| angptl3 | 114421 |
| angptl4 | 492647 |
| apoa1a | 30355 |
| apoa1b | 100101640 |
| apoa2 | 322327 |

| | |
|---|---|
| apoa4b.1 | 322543 |
| apoa4b.2 | 570354 |
| apoba | 566465 |
| apoc1 | 570638 |
| apoc2 | 568972 |
| apoea | 553587 |
| apoeb | 30314 |
| cetp | 492488 |
| cyp1a | 140634 |
| cyp1c1 | 553637 |
| cyp1d1 | 492344 |
| cyp20a1 | 406641 |
| cyp26a1 | 30381 |
| cyp26b1 | 324188 |
| cyp26c1 | 554036 |
| cyp27a7 | 402831 |
| cyp27c1 | 558396 |
| cyp2aa6 | 324212 |
| cyp2aa8 | 450060 |
| cyp2aa9 | 790957 |
| cyp2ad3 | 569245 |
| cyp2k16 | 449790 |
| cyp2k8 | 561462 |
| cyp2p10 | 399485 |
| cyp2v1 | 494153 |
| cyp2y3 | 368352 |
| cyp46a1.3 | 692332 |
| cyp4t8 | 387527 |
| hsd3b7 | 327462 |
| lcat | 793137 |
| ldlra | 387529 |
| ldlrap1a | 368278 |
| ldlrap1b | 791153 |
| lipg | 393096 |
| lpl | 30354 |
| lrp1aa | 563149 |
| lrp1ab | 565797 |
| lrp2a | 568184 |
| lrpap1 | 333939 |

| | |
|---|---|
| mylipa | 335888 |
| mylipb | 565911 |
| npc1 | 553330 |
| npc2 | 282673 |
| osbpl5 | 100006189 |
| pltp | 445125 |
| scarb1 | 387260 |
| sort1a | 406511 |
| sort1b | 799153 |
| stard3 | 63998 |
| sult1st1 | 323424 |
| sult2st1 | 338214 |
| sult4a1 | 678517 |
| sult6b1 | 322462 |
| sumo1 | 406438 |
| tspo | 450011 |
| vapal | 436819 |
| vapb | 323628 |
| vdac1 | 334582 |
| vdac2 | 322126 |
| vdac3 | 406529 |
| zgc:110366 | 550476 |
| hmgcra | 559054 |
| hmgcrb | 541479 |
| acat1 | 445290 |
| acat2 | 30643 |
| srebf2 | 100037309 |
| abcg1 | 556979 |

**Supplementary Table 4: Top 25 differentially expressed genes (DEG) in the cholesterol gene set in the head tissue of *tpp1*$^{-/-}$ mutant zebrafish compared with WT siblings.** Upregulated genes are in red and downregulated genes are in blue. Genes are ranked by their F value, showing most DEG in decreasing order from top to bottom. *Also in top 25 DEG in tail tissue of *tpp1*$^{-/-}$ mutant zebrafish compared with WT siblings (Supplementary Table 5).

| Gene | Protein | Human ortholog | Function | Reference |
|---|---|---|---|---|
| *apoeb** | Apolipoprotein Eb | *APOE* | major transporter of extracellular cholesterol in the brain | 1 |
| *cyp24a1** | Cytochrome P450, family 24, subfamily A, polypeptide 1 | *CYP24A1* | catabolism of vitamin D | 2 |
| *abca1a** | ATP-binding cassette, sub-family A (ABC1), member 1A | *ABCA1* | cholesterol exporter | 3 |
| *cyp27c1** | Cytochrome P450, family 27, subfamily C, polypeptide 1 | *CYP27C1* | unknown, retinoid 3,4-desaturase | 4 |
| *hsd11b2** | Hydroxysteroid (11-beta) dehydrogenase 2 | *HSD11B2* | conversion of cortisol to cortisone | 5 |
| *lrp1ab** | Low density lipoprotein receptor-related protein 1Ab | *LRP1* | member of LDL receptor family | 6 |
| *lpl** | Lipoprotein lipase | *LPL* | triglyceride hydrolase, regulates serum HDL levels | 7 |
| *lipf* | Lipase F, gastric type | *LIPF* | digestion of dietary triglycerides | 8 |
| *abca1b* | ATP-binding cassette, sub-family A (ABC1), member 1B | *ABCA1* | cholesterol exporter | 3 |
| *ugt2a4* | UDP glucuronosyltransferase 2 family, polypeptide A4 | | catalyses conjugation of lipophilic substrates, including hormones, with glucuronic acid to increase water solubility and enhance excretion | 9 |
| *vdac1** | Voltage-dependent anion channel 1 | *VDAC1* | gatekeeper for the passages of metabolites, nucleotides, and ions into the mitochondria | 10 |
| *lcat** | Lecithin-Cholesterol Acyltransferase | *LCAT* | esterifies free cholesterol | 3 |
| *angptl4* | Angiopoietin-like 4 | *ANGPTL4* | key inhibitory regulator of lipoprotein lipase | 11 |
| *sult6b1* | Sulfotransferase Family 6B Member 1 | *SULT6B1* | cytosolic sulfotransferase, possible role in thyroxine metabolism | 12 |
| *cyp4t8* | Cytochrome P450, family 4, subfamily T, polypeptide 8 | *CYP4B1* | ER cytochrome P450 enzyme | 13 |

| | | | | |
|---|---|---|---|---|
| *ldlra** | Low density lipoprotein receptor a | *LDLR* | Import of LDL | 3 |
| *srebf2** | Sterol regulatory element binding transcription factor 2 | *SREBF2/ SREBP2* | master transcriptional regulator of cholesterol biosynthesis | 3 |
| *cyp3c1* | Cytochrome P450, family 3, subfamily c, polypeptide 1 | *CYP3A4* | conversion of estrone to 16α-OH-estrone | 5 |
| *cyp11c1* | Cytochrome P450, family 11, subfamily C, polypeptide 1 | *CYP11B1* | conversion of testosterone to 11-ketotestosterone and of 11-deoxycortisol to cortisol | 5 |
| *sult2st1* | Sulfotransferase family 2, cytosolic sulfotransferase 1 | *SULT2A1/SULT2B1* | conversion of pregnenolone and DHEA to sulfated forms | 5 |
| *npc2* | NPC Intracellular Cholesterol Transporter 2 | *NPC2* | Lysosomal cholesterol export | 3 |
| *comta* | Catechol-O-methyltransferase a | *COMT* | O-methylation of endogenous neurotransmitters and hormones incorporating catecholic structures | 14 |
| *cyp26c1** | Cytochrome P450 Family 26 Subfamily C Member 1 | *CYP26C1* | retinoic acid metabolism | 15 |
| *hsd20b2* | Hydroxysteroid (20-beta) dehydrogenase 2 | | catalyses 20β-reduction of glucocorticoids | 16 |
| *cyp2aa6* | Cytochrome P450, family 2, subfamily AA, polypeptide 6 | | unknown | |

**Supplementary Table 5: Top 25 differentially expressed genes (DEG) in the cholesterol gene set in the tail tissue of *tpp1*$^{-/-}$ mutant zebrafish compared with WT siblings.** Upregulated genes are in red and downregulated genes are in blue. Genes are ranked by their F value, showing most DEG in decreasing order from top to bottom. *Also in top 25 DEG in head tissue of *tpp1*$^{-/-}$ mutant zebrafish compared with WT siblings (Supplementary Table 4).

| Gene | Protein | Human ortholog | Function | Reference |
|---|---|---|---|---|
| *apoeb** | Apolipoprotein Eb | *APOE* | major transporter of extracellular cholesterol in the brain | 1 |
| *cyp24a1** | Cytochrome P450, family 24, subfamily A, polypeptide 1 | *CYP24A1* | catabolism of vitamin D | 2 |
| *hsd11b2** | Hydroxysteroid (11-beta) dehydrogenase 2 | *HSD11B2* | conversion of cortisol to cortisone | 5 |
| *cyp26b1** | Cytochrome P450 Family 26 Subfamily C Member 1 | *CYP26C1* | retinoic acid metabolism | 15 |
| *lpl** | Lipoprotein lipase | *LPL* | triglyceride hydrolase, regulates serum HDL levels | 7 |
| *abca1a** | ATP-binding cassette, sub-family A (ABC1), member 1A | *ABCA1* | cholesterol exporter | 3 |
| *ldlra** | Low density lipoprotein receptor a | *LDLR* | Import of LDL | 3 |
| *cyp27c1** | Cytochrome P450, family 27, subfamily C, polypeptide 1 | *CYP27C1* | unknown, retinoid 3,4-desaturase | 4 |
| *cyp26c1** | Cytochrome P450 Family 26 Subfamily C Member 1 | *CYP26C1* | retinoic acid metabolism | 15 |
| *srebf2** | Sterol regulatory element binding transcription factor 2 | *SREBF2/ SREBP2* | master transcriptional regulator of cholesterol biosynthesis | 3 |
| *cyp2aa9* | Cytochrome P450, family 2, subfamily AA, polypeptide 9 | | prostaglandin metabolism | 17 |
| *tspo* | Translocator protein | *TSPO* | cholesterol import into the inner mitochondrial membrane | 18 |
| *hsd17b12a* | Hydroxysteroid (17-beta) dehydrogenase 12a | *HSD17B12* | conversion of 17β-Estradiol to estrone/estriol | 5 |
| *lipg* | Lipase G, Endothelial Type | *LIPG* | HDL metabolism | 19 |
| *lcat** | Lecithin-Cholesterol Acyltransferase | *LCAT* | esterifies free cholesterol | 3 |

| | | | | |
|---|---|---|---|---|
| *cyp2ad3* | Cytochrome P450, family 2, subfamily AD, polypeptide 3 | | unknown | |
| *apoc1* | Apolipoprotein C-I | *APOC1* | cholesterol catabolism | 20 |
| *mylipa* | Myosin regulatory light chain interacting protein a | *MYLIP* | regulates LDLR | 21 |
| *sult4a1* | Sulfotransferase family 4A, member 1 | *SULT4A1* | Cytosolic sulfotransferase | 22 |
| *cetp* | Cholesteryl ester transfer protein, plasma | *CETP* | HDL metabolism | 23 |
| *sult2st2* | Sulfotransferase family 2, cytosolic sulfotransferase 2 | | Cytosolic sulfotransferase | 24 |
| *vdac1** | Voltage-dependent anion channel 1 | *VDAC1* | gatekeeper for the passages of metabolites, nucleotides, and ions into the mitochondria | 10 |
| *stard3* | StAR-related lipid transfer (START) domain containing 3 | *STARD3* | mediates endoplasmic reticulum-to-endosome cholesterol transport at membrane contact sites | 25 |
| *lrp1ab** | Low density lipoprotein receptor-related protein 1Ab | *LRP1* | member of LDL receptor family | 6 |
| *lbr* | Lamin B receptor | *LBR* | cholesterol synthesis | 26 |
